# Arginine methylation promotes siRNA-binding specificity for a spermatogenesis-specific isoform of the Argonaute protein CSR-1

**DOI:** 10.1101/2020.06.08.134908

**Authors:** Dieu An H. Nguyen, Carolyn M. Phillips

**Author notes:** Corresponding author: Carolyn M. Phillips, Department of Biological Sciences, University of Southern California, 1050 Childs Way RRI 201, Los Angeles, CA 90089-2910.

## Abstract

CSR-1 is an essential Argonaute protein that binds to a subclass of 22G-RNAs targeting most germline-expressed genes. Here we show that the two isoforms of CSR-1 have distinct expression patterns; CSR-1B is ubiquitously expressed throughout the germline and during all stages of development while CSR-1A expression is restricted to germ cells undergoing spermatogenesis. Furthermore, CSR-1A associates preferentially with 22G-RNAs mapping to spermatogenesis-specific genes whereas CSR-1B-bound small RNAs map predominantly to oogenesis-specific genes. Interestingly, the exon unique to CSR-1A contains multiple dimethylarginine modifications, which are necessary for the preferential binding of CSR-1A to spermatogenesis-specific 22G-RNAs. Thus, we have discovered a regulatory mechanism for *C. elegans* Argonaute proteins that allows for specificity of small RNA binding between similar Argonaute proteins with overlapping temporal and spatial localization.

## Introduction

In the race for evolutionary fitness, sexual selection is a strong selective force. Multicellular eukaryotes have long favored sexual divergence of the male and female genders to increase genetic diversity, often resulting in sexual dimorphism. In *C. elegans*, a protandrous nematode, sexual reproduction can occur in a single germline where spermatogenesis and oogenesis take place sequentially. Both eggs and sperm are derived from the same pool of mitotically replicating nuclei at the distal tip of each gonad arm, which subsequently undergo meiosis as they progress through the germline toward the proximal end. During the last larval stage (L4), ~40 meiotic nuclei in each gonad arm begin the process of spermatogenesis, where they differentiate into mature sperm (Klass et al., 1976). Subsequently, oogenesis begins as the animal enters the adult stage, and the germline continues to produce oocytes for the rest of the reproductive cycle (Riddle et al., 1997).

Any given germ cell has the potential to differentiate into either spermatocytes or oocytes, though once committed to a sexual fate, the cells undergo distinct meiotic programs that vary remarkably in rate and mechanics (Shakes et al., 2009). Regardless of these differences, meiosis occurs exclusively in germ cells, which are surrounded by an environment inducive to the expression of germline-specific transcripts and suppression of somatic transcripts. One mechanism by which *C. elegans* germ cells promote this selective environment is by way of a perinuclear structure called the P granule (Knutson et al., 2017). These germline granules, which help to ensure pluripotency and germ cell identity, form a phase-separated condensate outside of the nucleus and contiguous with nuclear pores, where they regulate newly synthesized germline transcripts (Brangwynne et al., 2009; Knutson et al., 2017; Pitt et al., 2000; Sheth et al., 2010). One key regulator of P granule morphology and *C. elegans* germline development is the Argonaute protein, CSR-1 (<u>C</u>hromosome <u>S</u>egregation and <u>R</u>NAi deficient) (Claycomb et al., 2009; Updike and Strome, 2009).

Argonaute proteins are the core effectors of all small RNA-related pathways. These proteins bind small guide RNAs and regulate complementary transcripts by either directly cleaving target RNAs or recruiting cofactors that promote transcriptional or post-transcriptional gene regulation (Ghildiyal and Zamore, 2009; Hutvagner and Simard, 2008). Of the ~27 Argonaute proteins annotated in the *C. elegans* genome, only CSR-1 is essential for fertility, and it is reported to regulate over 4,000 germline-expressed genes (Claycomb et al., 2009; Yigit et al., 2006). CSR-1 binds to a class of antisense 22-nucleotide small interfering RNAs with a 5’ guanosine (22G-RNAs) whose target transcripts are thought to be protected from silencing by other small RNA-mediated pathways, and are thus “licensed” for germline expression (Claycomb et al., 2009; Seth et al., 2013; Wedeles et al., 2013). In contrast, foreign DNA such as transposons and transgenes containing non-*C. elegans* sequences are recognized by piwi-interacting RNAs (piRNAs, also known as 21U-RNAs in *C. elegans*), which, along with their Argonaute co-factor PRG-1, can initiate a multigenerational epigenetic silencing signal that depends on chromatin factors and 22G-RNAs bound by the worm-specific Argonaute proteins (WAGOs) (Ashe et al., 2012; Bagijn et al., 2012; Shirayama et al., 2012). The balance of gene licensing and gene silencing by CSR-1 and the WAGO proteins, respectively, is critical to promote expression of essential germline genes; and a disrupted balance severely compromises germline development and fertility (de Albuquerque et al., 2015; Phillips et al., 2015).

Studies of CSR-1 have reported several severe phenotypes that lead to defects in viability and fertility. In *csr-1* mutant embryos, chromosomes fails to properly align on the metaphase plate, resulting in aberrant mitotic division and anaphase bridging (Claycomb et al., 2009; Gerson-Gurwitz et al., 2016; Yigit et al., 2006). In the adult germline, loss of CSR-1 leads to fewer germ cells, defects in meiotic progression, and a delayed spermatogenesis to oogenesis switch (She et al., 2009). Additionally, CSR-1 has been implicated in a myriad of other cellular processes, including maturation of core histone mRNAs, promotion of sense-oriented RNA polymerase II transcription, attenuation of translation elongation, prevention of premature activation of embryonic transcripts in oocytes, alternative splicing, and paternal inheritance (Avgousti et al., 2012; Barberán-Soler et al., 2014; Cecere et al., 2014; Conine et al., 2010; Fassnacht et al., 2018; Friend et al., 2012). Yet, how one Argonaute protein can regulate so many seemingly distinct processes remains unanswered.

Here we demonstrate that CSR-1 has two isoforms – CSR-1B, which is present throughout the germline, and CSR-1A, which is specific to spermatogenic germ cells. CSR-1A and CSR-1B associate with distinct subsets of small RNAs to regulate spermatogenic or oogenic transcripts, respectively. The specificity of the two CSR-1 isoforms is interesting, considering they share nearly complete sequence homology and co-localize at the P granule in L4 larval and male germlines. We found that the first exon of CSR-1A is modified at arginine/glycine (RG) motifs by dimethylarginine. Here we show that loss of the dimethylarginine results in the loss of CSR-1A specificity for its preferred spermatogenic small RNA partners, resulting in CSR-1A indiscriminately binding to both spermatogenic and oogenic siRNAs. Thus, in this study, we have identified the first instance of methylarginine modification of a *C. elegans* Argonaute protein and demonstrated a previously unappreciated mechanism by which Argonaute proteins can acquire small RNA specificity.

## Results

### The long isoform of CSR-1 is selectively expressed during spermatogenesis

The Argonaute protein, CSR-1, has two isoforms, though previously little was known about their differential expression or function. The longer isoform, referred to as CSR-1A, and the shorter isoform, CSR-1B, share complete sequence homology, except for a unique exon at the 5’ end of CSR-1A (Figure 1A). Both isoforms are expressed, though at differential levels (Figure 1B) (Claycomb et al., 2009; Gerson-Gurwitz et al., 2016). To explore the distinct functions of the two CSR-1 isoforms, we used CRISPR/Cas9 to endogenously tagged CSR-1A at its N-terminus with a 2xHA/mCherry tag. We also tagged both CSR-1A and CSR-1B in the same strain by adding a 3xFLAG/GFP tag to the N-terminus of CSR-1B (Figure 1A). When both isoforms are tagged, CSR-1A+B is strongly expressed throughout the germline and during all developmental stages, in agreement with previous studies (Figure 1A) (Claycomb et al., 2009). Interestingly, when only CSR-1A is tagged, expression is restricted to the spermatogenesis region of the germline during the fourth larval stage (L4) of hermaphrodites and in males (Figure 1A and S1). We confirmed this observation by western blot analysis of all larval stage and adult animals, detecting CSR-1A expression robustly during the L4 stage and very weakly in gravid adults, coinciding temporally with spermatogenesis in *C. elegans* (Figure 1C). We next examined the differences in *csr-1a* and *csr-1b* transcript levels by performing RT-qPCR on wild-type animals using primers that recognize either *csr-1a* uniquely, or *csr-1a* and *csr-1b* together. We observed that the *csr-1a* transcript level is strongly upregulated during the L4 larval stage and then is reduced in gravid adults (Figure 1D). In contrast, using primers that recognize both *csr-1a* and *csr-1b*, we observed relatively stable transcript levels throughout development (Figure 1D). Taken together, these results demonstrate that though both CSR-1 isoforms are expressed in the germline, CSR-1A expression is limited to germ cells undergoing spermatogenesis while CSR-1B is expressed throughout the germline in larval and adult animals.

**Figure 1.**
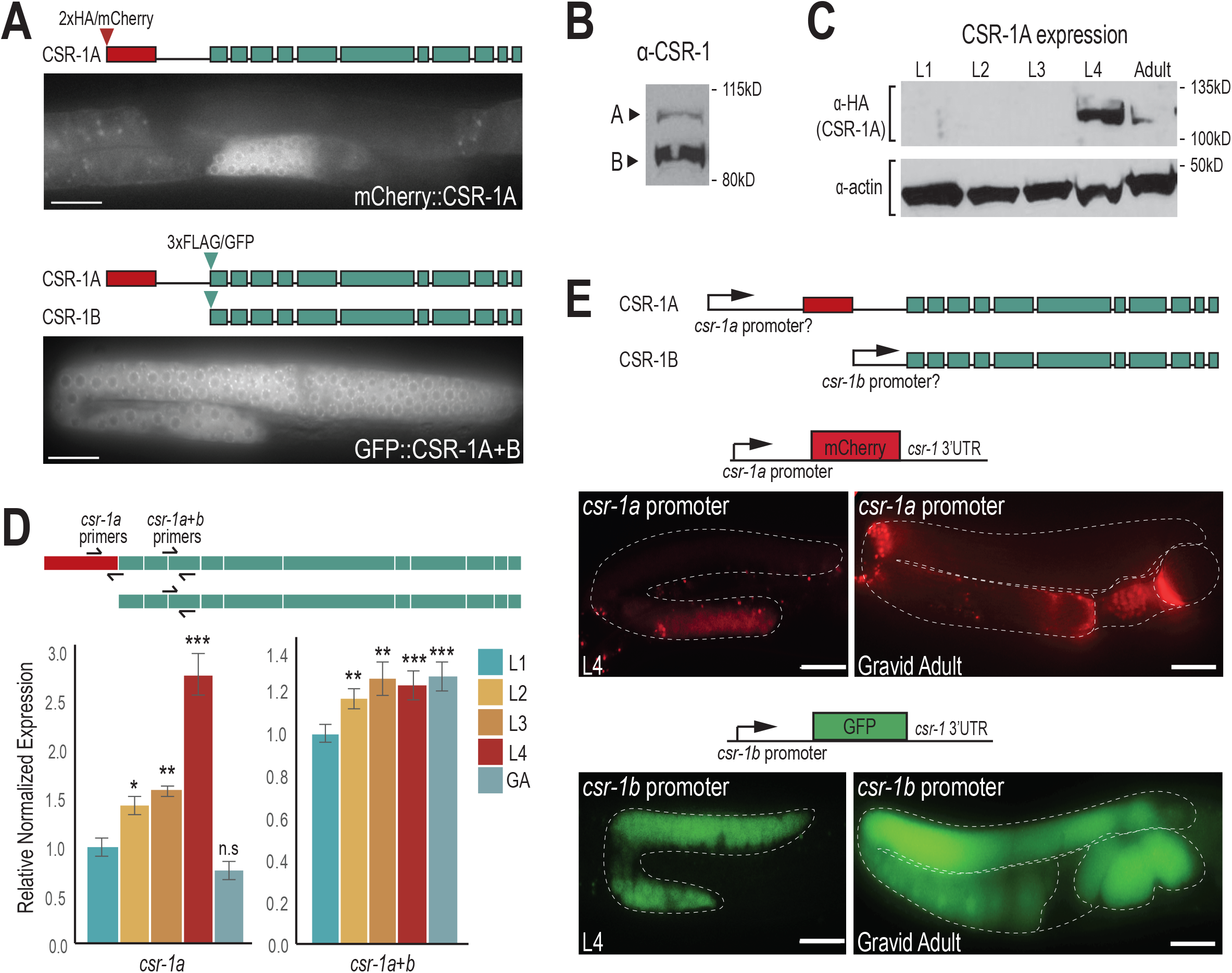
CSR-1 isoforms have distinct spatial and temporal expression patterns. (A) Live imaging of CSR-1A (top) and CSR-1A+B (bottom) in an L4 hermaphrodite germline. Scale bars, 25μM. (B) Western blot detecting CSR-1 expression, using α-CSR-1 antibodies, in wild-type animals at L4 stage. (C) Western blot detecting for CSR-1A at each larval stage and gravid adult in 2xHA/mCherry::CSR-1A strain using α-HA antibody. Actin is shown as a loading control. (D) Schematic (top) shows *csr-1* and qRT-PCR primer design. For qRT-PCR (bottom), RNA was isolated from wild-type animals at each larval stage and adult. Relative expression was normalized to *rpl-32* and Y45F10D.4, and calculated relative to L1 stage. Statistical significance was calculated for each stage compared to L1. * indicates a p-value≤0.05, ** indicates a p-value≤0.01, and *** indicates a p-value≤0.001. (E) Schematic representation (top) of putative *csr-1a* and *csr-1b* promoters driving mCherry and GFP. The *csr-1a* promoter comprises ~1.5kb of sequence preceding the first *csr-1a* exon, and the *csr-1b* promoter is the entire 544-bp intron between the unique *csr-1a* exon and the start codon of *csr-1b*. Schematic of the promoter reporter constructs created using the MoSCI system for *csr-1a* (middle) and *csr-1b* (bottom) are shown above images of gonads from L4 and gravid adult stages expressing the reporter constructs. Scale bars, 25μM.

### CSR-1A and CSR-1B are expressed from independent promoters

To address how CSR-1A and CSR-1B have such distinct expression patterns, we sought to determine whether they are expressed from independent promoters. We first fused ~1.5kb of DNA preceding the *csr-1a* start codon to an mCherry reporter and exogenously expressed it in the germline using the MoSCI system (Figure 1E) (Frøkjaer-Jensen et al., 2008). The region preceding the *csr-1a* start codon, the putative *csr-1a* promoter, drives mCherry expression only during the L4 stage and, during this time, expression is restricted to the spermatogenesis region of the germline. In gravid adults, residual mCherry expression can be detected inside the spermatheca, but is completely absent from the rest of the germline (Figure 1E). To determine whether any regulatory sequences reside in the intron between the unique *csr-1a* exon and the start codon for *csr-1b*, we fused ~0.5kb of DNA from this intron to a GFP reporter and expressed it in the germline using the MoSCI system (Figure 1E). This region, the putative *csr-1b* promoter, drives GFP expression throughout the germline cytoplasm, in all developmental stages. The GFP expression can be observed in oocytes and fertilized embryos but is excluded from the spermatheca (Figure 1E). These promoter fusion experiments corroborate our western blot and RT-qPCR analysis and indicate that CSR-1A is expressed predominantly during spermatogenesis, hinting at a unique role for CSR-1A during this developmental time point. Furthermore, the promoter fusion transgenes were able to recapitulate the expression pattern of the isoform-specific translation fusion proteins (Figure 1A), indicating that the promoters are sufficient to drive the distinct expression patterns. Together, these data indicate that the intronic region that sits between the first *csr-1a* exon and the start codon for *csr-1b* contains the promoter sequence for CSR-1B, and that their distinct promoters independently establish the differential expression patterns of CSR-1A and CSR-1B.

### CSR-1A localizes to the P granules during spermatogenesis

To further characterize the expression pattern of CSR-1A, we immunostained for CSR-1A and DAPI-stained for germline nuclei in dissected L4 hermaphrodite gonads. As the germline progresses towards the proximal end, germ cells undergo early stages of meiosis up until pachytene in a non-sex-specific manner (Shakes et al., 2009). In post-pachytene, however, spermatocytes and oocytes commit to their respective sexual fates, and spermatocyte chromosomes exhibit an extensive condensation phase (Shakes et al., 2009). During L4, when spermatogenesis begins, CSR-1A expression is first observed in cells that are exiting pachytene, namely cells transitioning into diplotene and karyosome, indicating that CSR-1A is selectively present in spermatocytes (Figure 2A). This expression is reminiscent of the sperm-specific Argonaute protein ALG-3, which is also present in spermatocytes during spermatogenesis (Conine et al., 2010). To further investigate the role of CSR-1A during spermatogenesis, we carefully examined protein localization at different time points. As ALG-3 is another Argonaute protein that is expressed during spermatogenesis and is required for proper sperm development, we created a double-transgenic strain labeling both CSR-1A and ALG-3, using CRISPR. At early L4, or 45 hours post hatching, both CSR-1A and ALG-3 form perinuclear foci in the spermatogenesis region that colocalize with the P granule marker PGL-1 (Figures 2B-C). In contrast, CSR-1A+B forms perinuclear foci throughout the germline (Figure S2A). As the animals reach the L4 to adult transition, 52 hours post hatching, CSR-1A and ALG-3 still colocalize during early spermatogenesis. However, by the time germ cells reach the secondary spermatocyte stage, CSR-1A expression becomes undetectable, while ALG-3 expression persists (Figure 2B, lower panel, Figure S2B). PGL-1 similarly disappears from P granules in primary spermatocytes and is subsequently cleared from secondary spermatocytes (Amiri et al., 2001). Together, these data demonstrate that like ALG-3, CSR-1A localizes to P granules during spermatogenesis, but unlike ALG-3, CSR-1A expression is restricted to the early stages of spermatogenesis and is not found in secondary spermatocytes.

**Figure 2.**
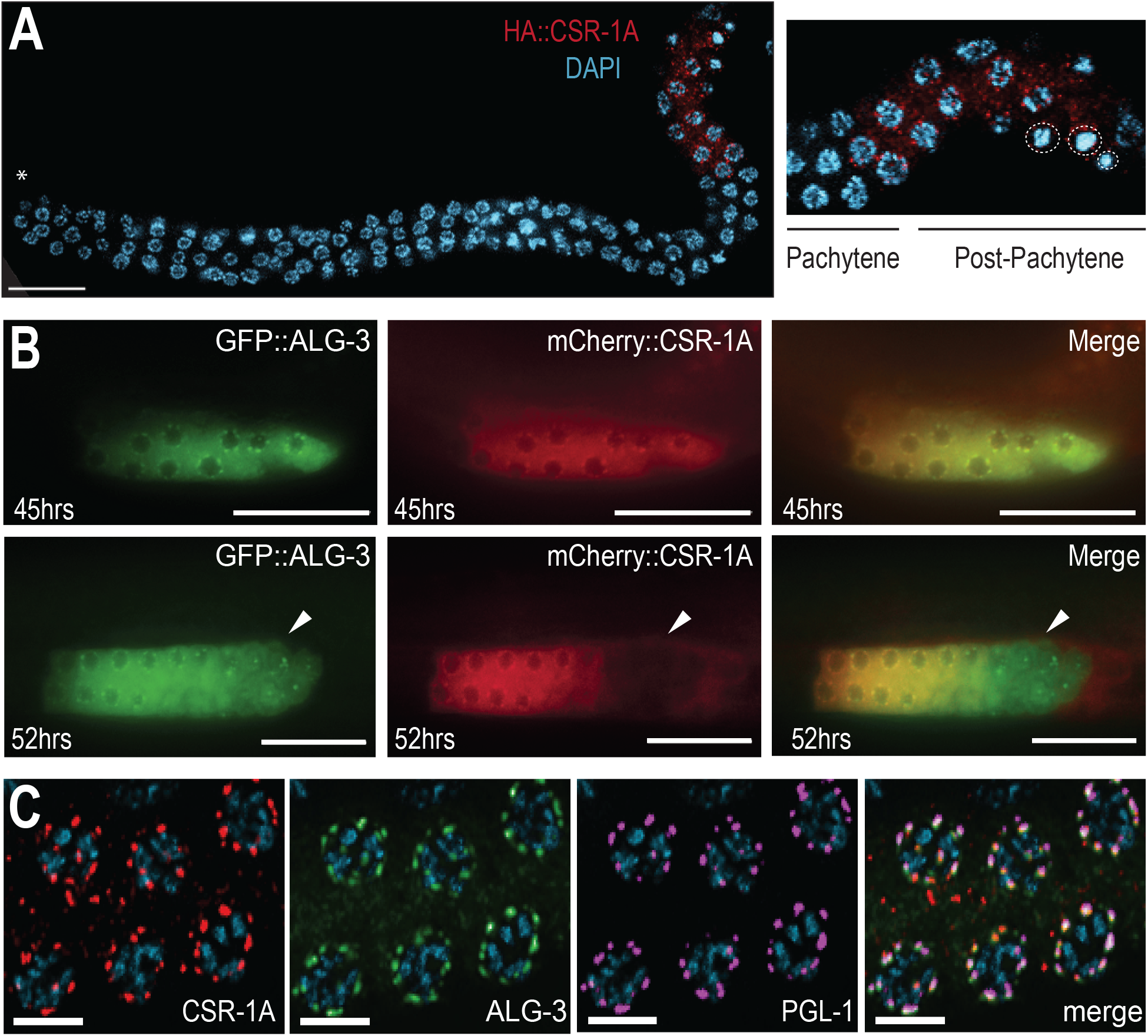
CSR-1A is expressed in the spermatogenesis region of the germline. (A) Immunofluorescence staining for HA::CSR-1A in dissected L4 hermaphrodite germline, using α-HA. Asterisk indicates distal end. Inset indicates region of cells exiting pachytene and entering post-pachytene where CSR-1A expression is detected. Dashed circles indicate karyosomes. DNA is stained with DAPI, blue. Scale bar, 15μM. (B) 3xFLAG/GFP::ALG-3 (left), 3xHA/mCherry::CSR-1A (middle), and merge (right) at 45 hrs (early L4 stage) or 52 hrs (young adult stage) post-L1 arrest. White arrows indicate region of secondary spermatocytes. Scale bars, 25 μM. (C) Immunofluorescent staining of a 2xHA::mCherry::CSR-1A; GFP::3xFLAG::ALG-3 dissected L4 hermaphrodite germline, using antibodies against HA, FLAG, and PGL-1. Scale bars represent 5μM.

### CSR-1A is required for optimal sperm fertility

We next sought to determine the role of CSR-1A in fertility. We first created two mutant alleles of *csr-1a* using CRISPR (Figure 3A). *csr-1a(cmp135)* removes the first seven amino acids including the start codon and *csr-1a(cmp143)* removes the same region plus ~500 bp of the *csr-1a* promoter region. We confirmed that the mutants have severely reduced *csr-1a* expression via RT-qPCR and western blot analysis (Figure S3A-B). We first examined the fertility of *csr-1a* mutants by comparing the brood size of *csr-1a*(*cmp135*) hermaphrodites to wild-type hermaphrodites at 25°C. We observed that *csr-1a*(*cmp135*) mutants had a modest but significant reduction in brood size compared to wild-type animals (Figure S3C). Because CSR-1A is expressed during spermatogenesis, we next sought to determine if CSR-1A is required for optimal male fertility. To this end, we assayed the ability of wild-type and *csr-1a* mutant male sperm to compete with wild-type hermaphroditic sperm by crossing males of the indicated genotypes to movement-defective, *unc-3*, hermaphrodites. We counted the number of progeny with the uncoordinated (Unc) phenotype, indicating self-progeny from hermaphrodite sperm, and with wild-type movement, indicating cross-progeny from male sperm. As expected, the wild-type male sperm outcompeted the hermaphroditic sperm for fertilization, due to their competitive advantages such as larger size and faster speed (Figure 3B) (LaMunyon and S. Ward, 1998; S. Ward and Carrel, 1979). However, when the *unc-3* hermaphrodites were crossed to *csr-1a* mutant males, the number of cross progeny was reduced significantly (~20%), indicating that *csr-1a* mutant male sperm are less successful at competing with hermaphroditic sperm (Figure 3B).

**Figure 3.**
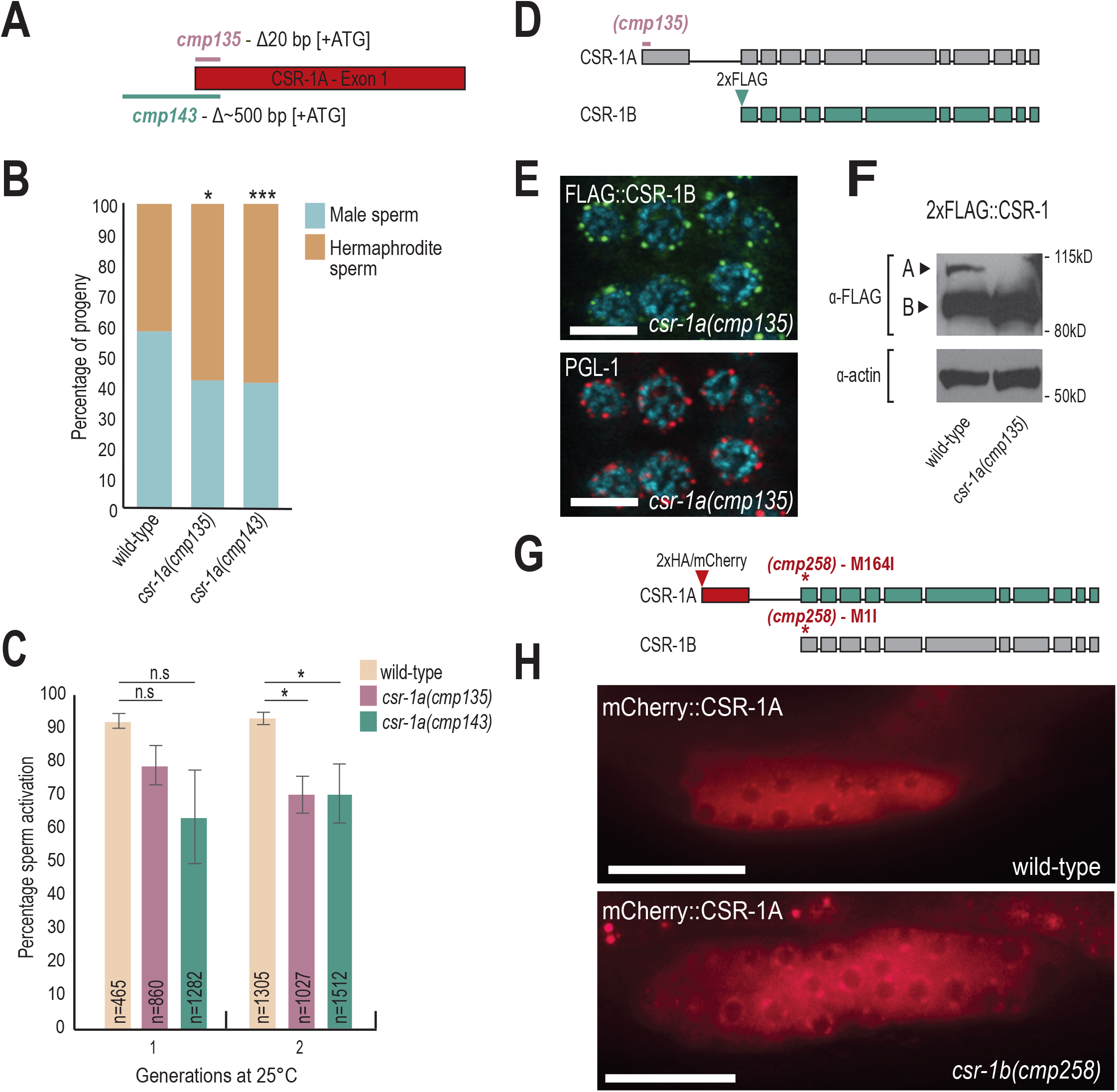
CSR-1 isoforms function independently of one another. (A) Schematic representation of two deletion alleles of *csr-1a*. *cmp135* removes 20 bp of coding sequence, including the start codon. *cmp143* removes ~500 bp of promoter sequence and 20 bp of coding sequence. (B) Male fertility assay performed on wild-type and mutant males crossed to *unc-3* hermaphrodite at 25°C. Unc progeny were scored as hermaphrodite sperm, and wild-type progeny were scored as male sperm. Two-tail T-test were performed to determine statistical significance. * indicates a p-value≤0.05 and *** indicates a p-value≤0.001. (C) An *in vitro* sperm activation assay was performed on spermatids from wild-type, *csr-1a(cmp135)* and *csr-1a(cmp143)* mutants. Animals were raised at 25°C for either one or two generations and each experiment was performed in triplicate. At least 100 spermatids were counted for each replicate for each condition, for a total of at least 400 spermatids per condition. Two-tail T-tests were performed to determine statistical significance. * indicates a p-value≤0.05 (D) Schematic representation of 2xFLAG::CSR-1B in a *csr-1a(cmp135)* mutant. The 2xFLAG tag was introduced immediately after the CSR-1B start codon in a *csr-1a(cmp135)* mutant animal. (E) Immunofluorescence staining for 2xFLAG::CSR-1B and P granules (PGL-1) in a *csr-1a(cmp135)* mutant at L4 stage. Scale bar, 5μM. (F) Western blot for CSR-1 expression levels, in wild-type and *csr-1a(cmp135)* mutant animals at L4 stage. Actin is shown as a loading control. (G) Schematic representation of disrupting *csr-1b* expression in a 2xHA/mCherry::CSR-1A strain. The *csr-1b* start codon was mutated from methionine to isoleucine (M1I) using CRISPR, which also introduces a M164I point mutation in the CSR-1A protein. (H) Live imaging of mCherry::CSR-1A in a wildtype and *csr-1b(cmp258)* mutant L4 hermaphrodite. Scale bar, 25 μM.

One aspect of sperm quality that can be tested *in vitro* is sperm activation. Sperm activation is the process by which round spermatids are transformed into mature and motile spermatozoa (Smith, 2014). This process can be recapitulated *in vitro* using Pronase E, a cocktail of proteases that induces sperm maturation and pseudopod formation (S. Ward et al., 1983). Because *C. elegans* fertility is reduced at 25°C, we raised wild-type and *csr-1a* mutant males at 25°C to assess sperm activation under compromised conditions. Wild-type spermatids from males that were raised at 25°C for one generation were activated at a rate of 92%, and after two generations at 25°C, they were activated at a rate of 93% (Figure 3C). In *csr-1a* males, we saw a modest reduction in the percentage of activated spermatids after one and two generations at 25°C (79% for *cmp135* and 63% for *cmp143* after one generation, and 63% for *cmp135* and 70% for *cmp143* after two generations) (Figure 3C). Because the effects we observed on sperm activation were modest yet significant, we generated two additional alleles of *csr-1a* to confirm our results. These new alleles remove the majority of the unique *csr-1a* exon (Figure S3D). Similar to what we observed with *csr-1a(cmp135)* and *csr-1a(cmp143)*, the two new alleles, *csr-1a(cmp253)* and *csr-1a(cmp254)* reduced the percentage of activated spermatids by approximately 20% (Figure S3E). Together, these data suggest that CSR-1A significantly contributes to optimizing male sperm quality, specifically when male sperm compete with hermaphroditic sperm for fertilization, in order to maximize male fitness.

### CSR-1A and CSR-1B are expressed independently of one another

Disruption of CSR-1 expression results in severe infertility and embryonic lethality (Claycomb et al., 2009; Yigit et al., 2006). To determine if our *csr-1a* alleles also affect the expression of CSR-1B protein, we introduced a 2xFLAG tag immediately following the start codon of CSR-1B in the *csr-1a*(*cmp135)* mutant background using CRISPR (Figure 3D). Similar to wild-type, CSR-1B in the *csr-1a*(*cmp135)* mutant localized to perinuclear foci associated with the P granule marker, PGL-1 (Figure 3E). Furthermore, by western blot, the CSR-1B protein was expressed at similar levels in wild-type and *csr-1a*(*cmp135)* mutant animals, whereas the CSR-1A was undetectable in the *csr-1a(cmp135)* mutant but present in the wild-type strain (Figure 3F). To determine if CSR-1B is required for CSR-1A expression, we generated an effectively null allele of *csr-1b*, where we mutated the *csr-1b* start codon to isoleucine, in the mCherry::CSR-1A strain using CRISPR (Figure 3G). In this *csr-1b(cmp258)* mutant, mCherry::CSR-1A is expressed at perinuclear foci in the spermatogenesis region of L4 animals, similar to wild-type expression of CSR-1A (Figure 3H). These animals, however, are sterile and need to be maintained over a balancer, a phenotype previously associated with *csr-1* mutants (Yigit et al., 2006). Together, these data demonstrate that the *csr-1a(cmp135)* mutant does not affect CSR-1B expression. More importantly, CSR-1A and CSR-1B do not depend on one another for localization or expression, and thus they behave as independent proteins with seemingly specialized functions in the *C. elegans* germline.

### CSR-1A and CSR-1B target distinct groups of genes

To investigate the small RNAs bound by each CSR-1 isoform and thus determine whether CSR-1A and CSR-1B have distinct small RNA partners, we immunoprecipitated CSR-1A (from the *2xHA::csr-1a* strain), CSR-1B (from the *csr-1a(cmp135) 2xFLAG::csr-1b* strain), and CSR-1A+B (from the *2xFLAG::csr-1b* strain), and sequenced the associated small RNAs. CSR-1A, CSR-1B, and CSR-1A+B were immunoprecipitated from L4 stage animals, and CSR-1B and CSR-1A+B were additionally immunoprecipitated from the adult stage, for comparison. We found that the small RNAs that immunoprecipitate with CSR-1A, CSR-1B, and CSR-1A+B are enriched for germline genes at both L4 and adult stages (Figure 4A and S4A). However, CSR-1A preferentially associates with small RNAs mapping to spermatogenic genes at the L4 stage while CSR-1B preferentially associates with small RNAs mapping to oogenic genes both at L4 and adult stages (Figure 4A and S4A). CSR-1B is expressed at much higher levels than CSR-1A at the L4 stage (Figure 1B and 3F), therefore we would expect that when we tag both isoforms together (CSR-1A+B) the majority of the small RNAs immunoprecipitated would associate with CSR-1B. Indeed, we observed that the majority of the small RNAs immunoprecipitated with CSR-1A+B map to oogenic genes at both L4 and adult stages (Figure 4A and S4A). Regardless of isoform, the majority of the small RNAs associated with CSR-1 are 22-nucleotide long with a strong bias for guanine as the 5’ nucleotide (Figure S4B-D). These data indicate that, while both isoforms of CSR-1 bind 22G-RNAs mapping to germline genes, CSR-1A is enriched for siRNAs mapping to spermatogenesis-specific genes and CSR-1B is enriched for siRNAs mapping to oogenesis-specific genes.

**Figure 4.**
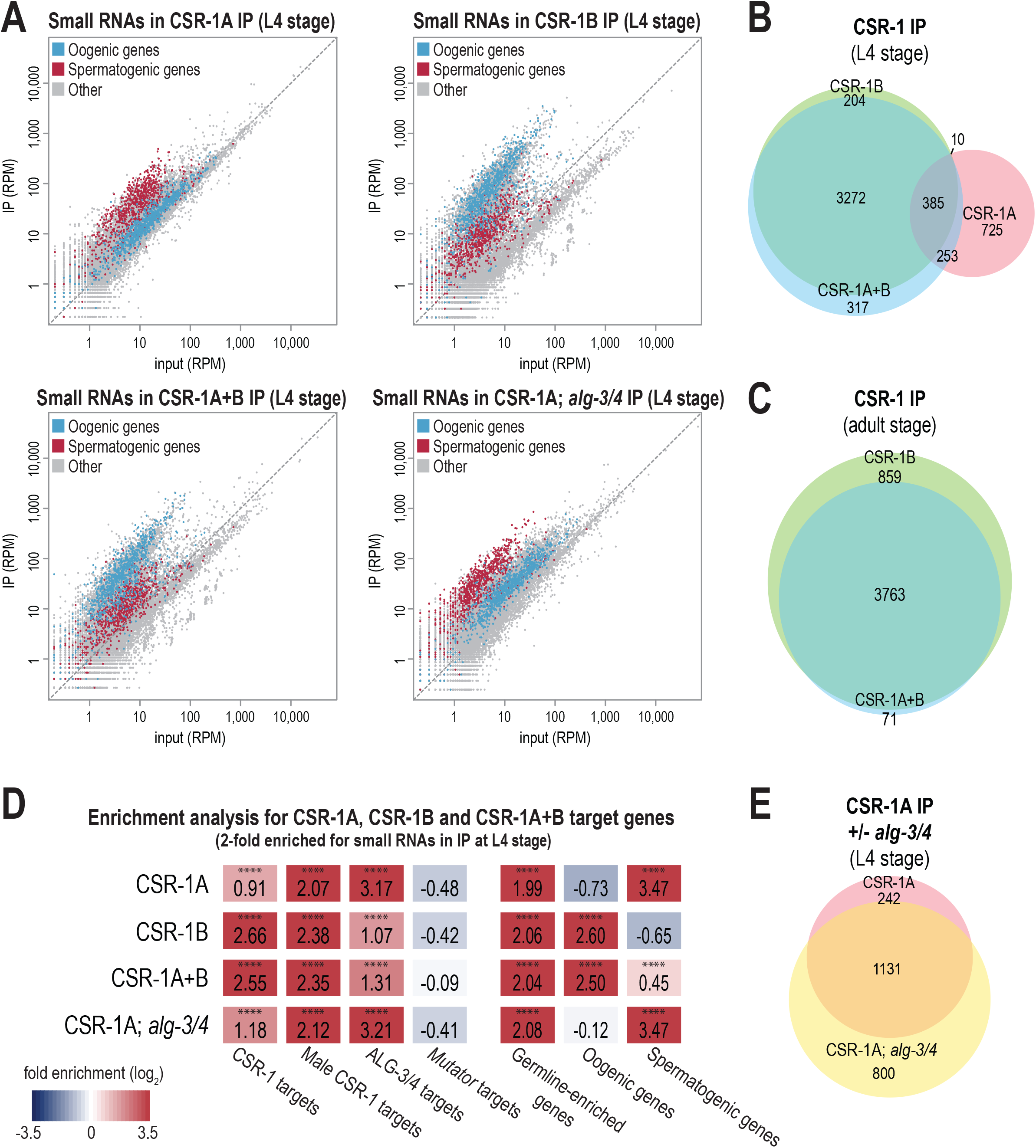
CSR-1A and CSR-1B target distinct groups of genes. (A) Normalized reads for oogenesis and spermatogenesis genes at L4 stage from HA::CSR-1A IP, FLAG::CSR-1B IP, FLAG::CSR-1A+B IP, and HA::CSR-1A; *alg-3/4* IP compared to input. Oogenesis and spermatogenesis genes are indicated in blue and red respectively. (B) Venn diagram indicates overlap of CSR-1A, CSR-1B, and CSR-1A+B target genes at L4 stage. Target genes are defined are at least 2-fold enriched in the IP, with at least 10 RPM in IP samples and a DESeq2 adjusted p-value≤0.05. (C) Venn diagram indicates overlap of CSR-1B and CSR-1A+B gene lists at adult stage. (D) Enrichment analysis (log2(fold enrichment)) examining the overlap of CSR-1A, CSR-1B, and CSR-1A+B target genes with known targets of the CSR-1, male CSR-1, ALG-3/4, and *mutator* small RNA pathways and oogenesis and spermatogenesis-enriched genes. See Materials and Methods for gene list information. Statistical significance for enrichment was calculated using the Fisher’s Exact Test function in R. **** indicates a p-value≤0.0001. (E) Venn diagram indicates overlap of CSR-1A target genes in the presence or absence of *alg-4; alg-3.*

We next sought to define a list of genes targeted by CSR-1A and CSR-1B. We defined the CSR-1A, CSR-1B, and CSR-1A+B target genes at both L4 and adult stages as those with complementary small RNAs at least two-fold enriched in the IP compared to input, with at least 10 RPM in the IP samples and a DESeq2 adjusted p-value of ≤0.05 (Table S1). Comparing these gene lists with one another, we found that the majority of CSR-1B target genes are also CSR-1A+B target genes at both L4 (94.5% overlap) and adult stages (81.4% overlap) (Figure 4B-C). Furthermore, the CSR-1B target genes at L4 stage overlap significantly with the CSR-1B target genes at adult stage (94.0% overlap), and similarly the CSR-1A+B target genes at L4 stage overlap significantly with the CSR-1A+B target genes at adult stage (78.0% overlap) (Figure S4E-F). In contrast, only 28.8% of CSR-1A target genes at L4 stage overlap with CSR-1B target genes at L4 stage, and only 46.6% of CSR-1A target genes at L4 stage overlap with CSR-1A+B target genes at L4 stage, despite the CSR-1A+B IP at L4 stage immunoprecipitating both isoforms (Figure 4B). Thus, these data demonstrate that CSR-1A targets a distinct set of genes and that due to the much higher expression of CSR-1B, even at L4 stage, immunoprecipitation of the two isoforms together enriches for small RNAs that predominantly map to CSR-1B target genes.

To further our understanding of which genes are targeted by CSR-1A and CSR-1B, we compared the CSR-1A, CSR-1B, and CSR-1A+B target gene lists to previously described target gene lists for other small RNA pathways (Conine et al., 2013; Gu et al., 2009; Lee et al., 2012; Phillips et al., 2014; Reinke et al., 2004; Tsai et al., 2015; Zhang et al., 2011). As expected, CSR-1B and CSR-1A+B are strongly enriched for hermaphrodite and male CSR-1 target genes, and more modestly enriched for ALG-3/4 target genes (Figure 4D). In contrast, CSR-1A is only modestly enriched for hermaphrodite CSR-1 target genes, and strongly enriched for ALG-3/4 target genes and male CSR-1 target genes (Figure 4D). It has previously been shown that male CSR-1 target genes, identified by immunoprecipitating both isoforms of CSR-1 in males, include both oogenesis-specific genes identified as CSR-1 targets in hermaphrodites, and male-specific genes that are also ALG-3/4 targets (Conine et al., 2013). Thus, it is not surprising that both CSR-1B, which preferentially binds hermaphrodite CSR-1 target genes, and CSR-1A, which preferentially binds ALG-3/4 target genes, are enriched for male CSR-1 target genes (Figure 4D). In further agreement with these data, the CSR-1A target gene list is strongly enriched for spermatogenic genes, the CSR-1B target gene list is strongly enriched for oogenic genes, and the CSR-1A+B gene list is strongly enriched for oogenic genes and more modestly enriched for spermatogenic genes (Figure 4D). In contrast, neither CSR-1A, CSR-1B, nor CSR-1A+B are enriched for *mutator* target genes, which include pseudogenes, repetitive elements, transposons, and other germline-repressed genes (Figure 4D). Together, these data further show that CSR-1A and CSR-1B bind small RNAs targeting germline genes associated with distinct small RNA pathways, with CSR-1A preferentially targeting spermatogenic genes and ALG-3/4 targets and CSR-1B preferentially targeting oogenic genes and hermaphrodite CSR-1 targets.

### CSR-1A does not require ALG-3/4 to bind spermatogenic small RNAs

Because CSR-1A shares a similar expression pattern to ALG-3 and targets spermatogenic genes, we next sought to determine whether the small RNAs bound to CSR-1A require ALG-3/4 for their production. To this end, we immunoprecipitated CSR-1A in an *alg-3/4* mutant background and sequenced the associated small RNAs. Interestingly, in the absence of *alg-3* and *alg-4*, CSR-1A was still preferentially loaded with small RNAs targeting spermatogenic genes (Figure 4A and 4D). In addition, 82.4% of CSR-1A target genes in the wild-type background were also defined as targets of CSR-1A in the *alg-3; alg-4* double mutant background (Figure 4E). Furthermore, CSR-1A target genes in the *alg-3; alg-4* double mutant background were strongly enriched for ALG-3/4 target genes and male CSR-1 target genes, and modestly enriched for hermaphrodite CSR-1 target genes, similar to CSR-1A target genes in a wild-type background (Figure 4D). Thus, these data show, despite CSR-1A and ALG-3/4 targeting many of the same genes, CSR-1A is not directly downstream of ALG-3/4 in regulating spermatogenic gene expression and ALG-3 and ALG-4 are dispensable for the production of CSR-1A-bound spermatogenic small RNAs.

### CSR-1A expression is positively correlated with the expression of CSR-1A target genes

It is well-established that CSR-1 targets germline-expressed genes, and that CSR-1 can either license and protect these transcripts from silencing by the piRNA pathway in the adult germline or can modestly tune the transcript level in developing oocytes and embryos (Claycomb et al., 2009; Gerson-Gurwitz et al., 2016; Seth et al., 2013). To determine whether CSR-1A is similarly protecting or tuning its target transcripts, we generated mRNA high-throughput sequencing libraries from wild-type and *csr-1a(cmp135)* mutants at 20°C and 25°C. At permissive temperature (20°C), CSR-1A targets are modestly, but significantly, downregulated in *csr-1a(cmp135)* mutants compared to wild-type (Figure 5A). Similarly, spermatogenic genes are modestly, but significantly, downregulated in *csr-1a(cmp135)* mutants compared to wild-type (Figure 5A). At elevated temperature (25°C), the reduced expression of CSR-1A target genes and spermatogenic genes in *csr-1a(cmp135)* mutants compared to wild-type becomes more substantial, however CSR-1B target genes and oogenic genes are also reduced in expression, consistent with a non-specific effect on germline-expressed genes at elevated temperature (Figure 5A). In contrast, *mutator* target genes were unchanged at both 20°C and 25°C in the *csr-1a(cmp135)* mutants (Figure 5A). To confirm that CSR-1A target transcripts are indeed reduced in the *csr-1a(cmp135)* mutant, we performed RT-qPCR on three spermatogenic genes that were strongly enriched for mapped small RNAs in the CSR-1A immunoprecipitation. We determined that all three genes showed a consistent and significant decrease in transcript level in the *csr-1a(cmp135)* mutant compared to wild-type animals (Figure 5B). Therefore, CSR-1A appears to have modest but significant positive effects on its target transcripts, such that in its absence, these genes have reduced expression.

**Figure 5.**
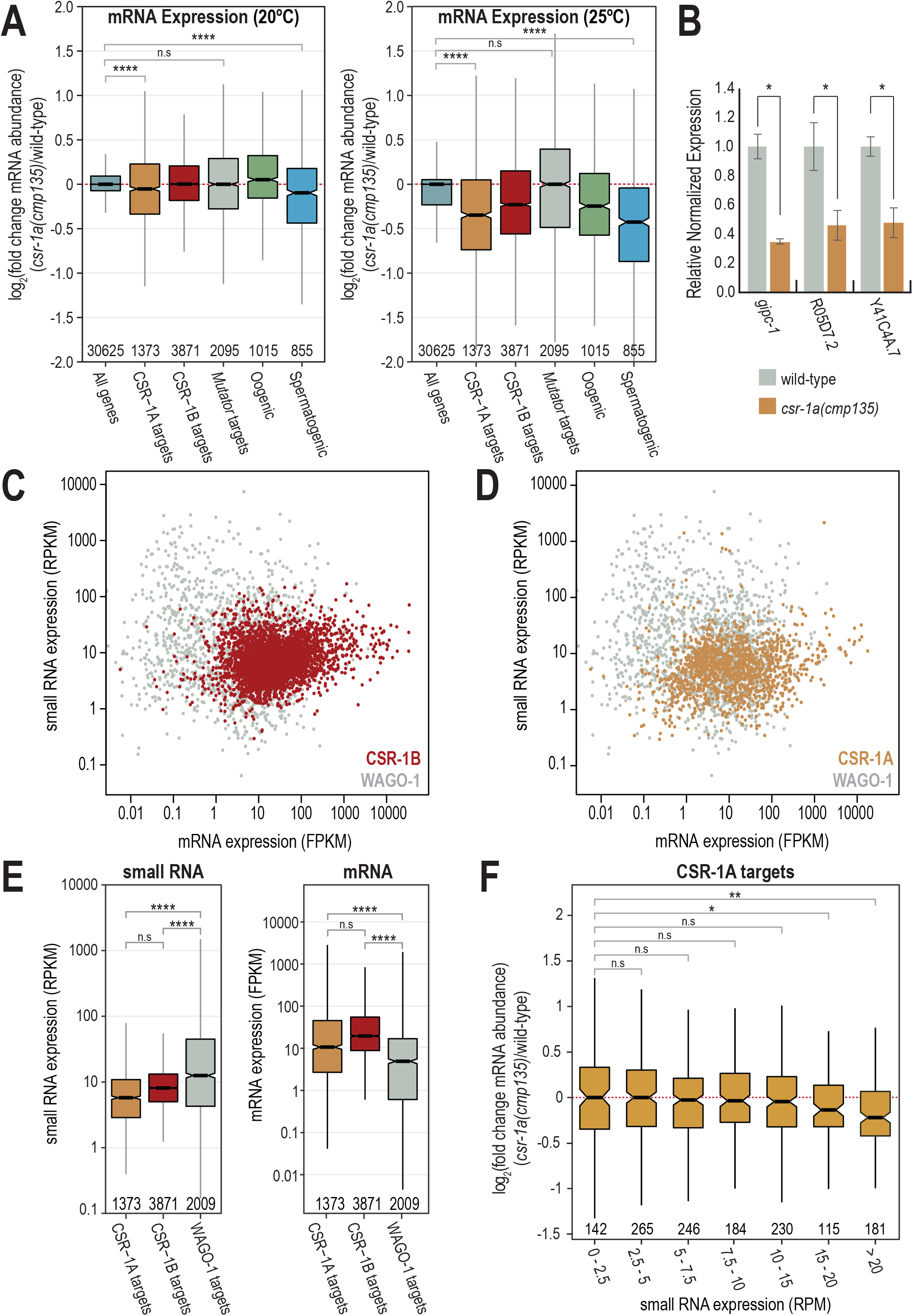
CSR-1A targets spermatogenesis-expressed genes. (A) Box plots depicting log2(fold change mRNA abundance) for L4 stage *csr-1a(cmp135)* mutants relative to wild-type animals at 20°C (left) and 25°C (right). Statistical significance was calculated in R using the Student’s t-test. n.s. denotes not significant and **** indicates a p-value≤0.0001. (B) RT-qPCR from wild-type and *csr-1a (cmp135)* L4 animals for *gipc-1*, R05D7.2, Y41C4A.7, genes that are highly enriched in HA::CSR-1A IP. Relative expression was normalized over *rpl-32*, and calculated relative to wild-type animals. * indicates a p-value≤0.05. (C-D) Small RNA expression (RPKM) plotted against mRNA expression (FPKM) in wild-type animals for CSR-1B and CSR-1A target genes compared against WAGO-1 target genes. CSR-1B, CSR-1A, and WAGO-1 gene lists come from FLAG::CSR-1B IP, HA::CSR-1A IP, and FLAG::WAGO-1 IP in L4 animals, respectively. (E) Box plot depicting small RNA reads per kilobase per million (RPKM, left) and mRNA fragments per kilobase per million (FPKM, right) from L4 wild-type animals, raised at 20°C, 48 hours post L1. Small RNAs and mRNAs transcripts were grouped based on their mapping to CSR-1B, CSR-1A, and WAGO-1 target genes, identified from the RNA-IP experiments. (F) Box plot depicting log2(fold change mRNA abundance) of CSR-1A target genes for L4 stage *csr-1a(cmp135)* mutants relative to wild-type animals at 20°C where the CSR-1A target genes are binned based on the mapped small RNA reads per million (RPM) in wild-type animals. Statistical significance was calculated using the Student’s t-test in R. n.s denotes not significant **** indicates a p-value≤0.0001.

Recent studies have examined the relationship between the levels of siRNAs targeting each transcript in correlation with mRNA expression (Bezler et al., 2019; Gerson-Gurwitz et al., 2016). In general, mRNAs expression tends to be lower for genes with very high levels of siRNAs and for genes that are targets of the Argonaute protein, WAGO-1 (Bezler et al., 2019). In contrast, mRNA expression tends to be higher for genes that are targets of CSR-1 (Bezler et al., 2019). To determine whether this trend holds true for CSR-1A target genes, for comparison, we generated a list of WAGO-1 target genes at the L4 stage by immunoprecipitating WAGO-1 and defining targets as those at least two-fold enriched in a WAGO-1 IP compared to input, with at least 10 RPM in the IP samples, and a DESeq2 adjusted p-value of ≤0.05. Then using mRNA and small RNA sequencing libraries from wild-type animals at L4 stage, we found that WAGO-1 targets have significantly higher levels of mapping complementary small RNAs compared to CSR-1A and CSR-1B targets (Figure 5C-E). Moreover, mRNA expression of WAGO-1 targets is significantly lower than mRNA expression of CSR-1A and CSR-1B target genes (Figure 5C-E). We further grouped the CSR-A target genes into seven bins based on their small RNA expression in wild-type animals. We found that higher levels of CSR-1A siRNAs (>15 RPM in wild-type animals) correlates with reduced mRNA expression in *csr-1a* mutants, while the expression of genes with lower levels of CSR-1A siRNAs tends to be unchanged in *csr-1a* mutants (Figure 5F). This result is consistent with both the very modest effect of the *csr-1a(cmp135)* mutant on transcript level of all CSR-1A target genes (Figure 5A) and the much more significant effect of the *csr-1a(cmp135)* mutant on the transcript level of the three CSR-1A target genes tested by qPCR (Figure 5F), which were selected based on the strong enrichment for their mapped small RNAs in the CSR-1A immunoprecipitation. Together, these data suggest that CSR-1A may share a similar licensing mechanism to what has been previously proposed for CSR-1, and that in *csr-1a* mutants, the spermatogenic transcripts licensed by CSR-1A are no longer protected and may be subjected to degradation.

### The first exon of CSR-1A is unstructured and contains RG motifs

CSR-1A differs from its shorter counterpart by a single N-terminal exon. The exon is arginine/glycine-rich (containing RG motifs), and we sought to determine whether this unique exon is conserved across the *Caenorhabditis* genus. We first identified CSR-1 orthologs in several closely related nematode species, including *C. brenneri*, *C. briggsae*, *C. japonica*, *C. latens*, *C. nigoni*, *C. remanei*, *C. sinica*, and *C. tropicalis*. The first exon of CSR-1 in each species is the least conserved portion of the protein, however, like *C. elegans* CSR-1A, each ortholog had enrichment of arginines and glycines in this region (Figure S5). Furthermore, while only a single isoform is annotated in most of the species, all orthologs possessed a conserved methionine at the position of the *C. elegans* CSR-1B start codon (Figure S5). Additionally, all eight orthologs have a large intron between the first exon and the rest of the protein, with a median size of 617 bp and the smallest being 503 bp in *C. tropicalis*. These data suggest that the regulatory elements driving CSR-1B expression in *C. elegans*, which are found in this intron, may also be conserved.

Because of the repetitive nature of the RG motif region and the lack of otherwise strong sequence conservation, we next asked whether the first exon of CSR-1 contains regions of intrinsic disorder. Using IUPred2, we determined that the first exon of CSR-1A in *C. elegans* is highly disordered while the rest of the protein is predicted to be structured (Figure S6A). We then examined the CSR-1 orthologs in related *Caenorhabditis* species and found that, like in *C. elegans*, the first exon of CSR-1 is highly disordered in all related nematode species (Figure S6B), demonstrating that the disordered nature of the first exon is a conserved feature of the CSR-1 protein.

### The RG motifs in the first exon of CSR-1A are dimethylated

It has been shown previously that RG motifs are often targets of Protein Arginine Methyltransferases (PRMTs), which catalyze the methylation of arginine residues (Kirino et al., 2009; 2010; Nishida et al., 2009; Vagin et al., 2009). To determine whether CSR-1A contains methylated arginines, we immunoprecipitated 2xHA::CSR-1A and subjected the sample to mass spectrometry analysis. We identified six dimethylated arginines in the first exon of CSR-1A, which constituted 100% of the RG motifs captured by mass spectrometry (Figure 6A and Table S2). The remaining RG motifs were not captured, methylated or unmethylated, by our mass spectrometry experiment. We did not identify any dimethylarginines in the portion of the CSR-1 protein found in both CSR-1A and CSR-1B isoforms, however the conserved RGRG sequence found near the N terminus of CSR-1B, a good candidate region for dimethylation, was also not captured by our mass spectrometry experiment. Together, these data indicate that the first exon of CSR-1A is heavily methylated, and, given the conservation of RG motifs in this region across *Caenorhabditis* species, this methylation is likely also conserved.

**Figure 6.**
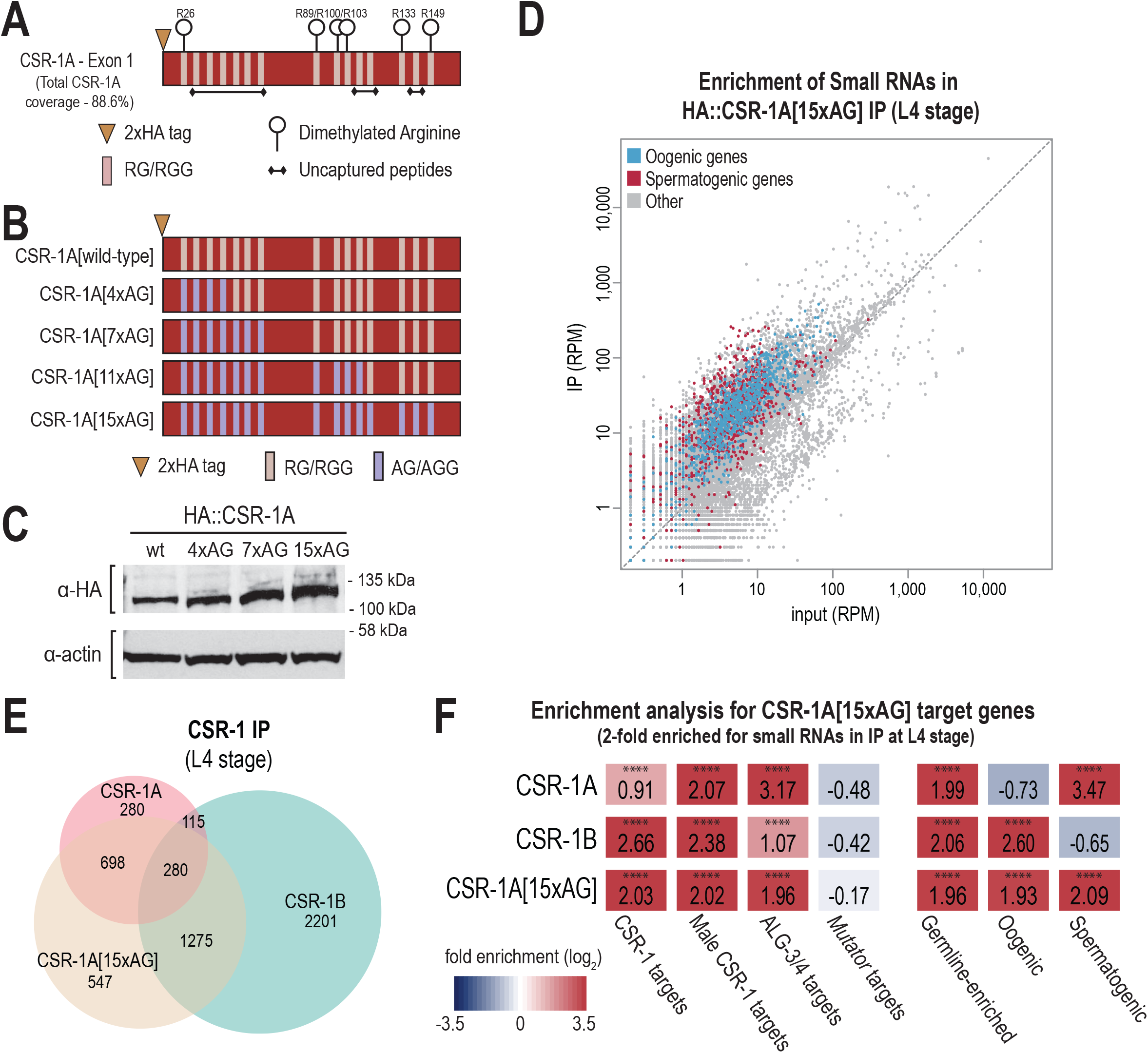
Methylation of CSR-1A N-terminal exon promotes binding preference for spermatogenic siRNAs. (A) Graphical display of dimethylation modifications detected on CSR-1A by mass spectrometry following IP. (B) Schematic representation of the series of arginine to alanine mutations created by CRISPR to generate the unmethylatable CSR-1A[15xAG]. (C) Western blot for HA::CSR-1A in the RG-to-AG mutants. Actin is shown as a loading control. (D) Normalized reads for spermatogenesis and oogenesis genes from HA::CSR-1A[15xAG] IP compared to input. Spermatogenesis and oogenesis are highlighted in red and blue respectively. (E) Venn diagram indicates overlap of CSR-1A, CSR-1B, and CSR-1A[15xAG] target genes at L4 stage. (F) Enrichment analysis (log2(fold enrichment)) examining the overlap of CSR-1A, CSR-1B, and CSR-1A[15xAG] target genes with known targets of the CSR-1, male CSR-1, ALG-3/4, and *mutator* small RNA pathways and oogenesis and spermatogenesis-enriched genes. See Materials and Methods for gene list information. Statistical significance for enrichment was calculated using the Fisher’s Exact Test function in R. **** indicates a p-value≤0.0001.

We next sought to study the function of the dimethylarginine modification. To this end, we mutated all arginines found in RG motifs within the first exon of CSR-1A to alanine using CRISPR, thus rendering the exon unmethylatable. Because we generated the mutations sequentially from the N-terminus of the protein, three-to-four arginines at a time, we created a series of proteins with four, seven, 11 or all 15 RG motifs mutated to AG – hereafter referred to as CSR-1A[4xAG], CSR-1A[7xAG], CSR-1A[11xAG] and CSR-1A[15xAG] (Figure 6B). We first asked if dimethylation of arginines affects the expression or stability of the CSR-1A protein. By western blot, we determined that the CSR-1A[4xAG], CSR-1A[7xAG] and CSR-1A[15xAG] proteins are expressed at levels comparable to wild-type (Figure 6C). Because dimethylation has been shown previously to contribute to Argonaute protein localization (Webster et al., 2015), we next tested whether dimethylation is necessary for CSR-1A localization to P granules. We performed immunostaining on CSR-1A[wild-type], CSR-1A[4xAG], and CSR-1A[15xAG] L4 stage hermaphrodites, and detected no discernable differences in localization in the RG-to-AG mutant strains (Figure S6C). Finally, to determine whether the demethylated RG motifs are required for fertility, we performed brood size assays on CSR-1A[wild-type], CSR-1A[7xAG], and CSR-1A[15xAG] animals. We found that CSR-1A[7xAG] and CSR-1A[15xAG] animals have a modest but significant reduction in brood size compared to the control strain, indicating that *csr-1a* RG motif mutants phenocopy a null *csr-1a* mutant (Figure S6D). Therefore, our data demonstrates that the first exon of CSR-1A contains the dimethylarginine modification, however this modification is not required for expression or localization of the CSR-1A protein.

### The RG motifs in CSR-1A promote specificity for binding small RNAs targeting spermatogenic genes

CSR-1A and CSR-1B both localize to P granules in the spermatogenesis region of the germline in L4 stage hermaphrodites and males, yet each isoform has distinct small RNA targets (Figure 1A and 4A-B). Because the isoforms only differ by the presence of the RG-motif-containing exon in CSR-1A, we asked if the RG-motifs and the dimethylarginine modification could be a mechanism by which CSR-1A preferentially associates with spermatogenic small RNAs. To test this hypothesis, we immunoprecipitated 2xHA::CSR-1A[15xAG] from L4 stage animals and subjected the associated small RNAs to high-throughput sequencing. We found that the small RNAs that immunoprecipitate with CSR-1A[15xAG] map to both spermatogenic and oogenic genes, with no discernable preference for either (Figure 6D). We next defined the list of genes targeted by CSR-1A[15xAG] as those at least two-fold enriched in the IP compared to input, with at least 10 RPM in the IP samples and a DESeq2 adjusted p-value of ≤0.05 (Table S1). Comparing this gene list to the wild-type CSR-1A and CSR-1B target gene lists at L4 stage, we found that 34.9% (978 genes) of the genes in the CSR-1A[15xAG] list overlapped with the wild-type CSR-1A gene list and 55.5% (1555 genes) of the genes in the CSR-1A[15xAG] list overlapped with the CSR-1B gene list (Figure 6E). Furthermore, when we compared the CSR-1A[15xAG] target gene list to previously described target genes lists for other small RNA pathways (Conine et al., 2013; Gu et al., 2009; Lee et al., 2012; Phillips et al., 2014; Reinke et al., 2004; Tsai et al., 2015; Zhang et al., 2011), we found that the CSR-1A[15xAG] target gene list is strongly enriched for hermaphrodite and male CSR-1 target genes, and for ALG-3/4 target genes (Figure 6F). Additionally, the CSR-1A[15xAG] target gene list is strongly enriched for both spermatogenic and oogenic genes (Figure 6F). These results are in contrast to the wild-type CSR-1A target gene list, which is strongly enriched for spermatogenic genes but not oogenic genes, and to the CSR-1B target gene list, which is strongly enriched for oogenic genes but not spermatogenic genes (Figure 6F). However, like wild-type CSR-1A and CSR-1B, CSR-1A[15xAG] is not enriched for *mutator* target genes (Figure 6F). Together, these data indicate that loss of the RG motifs, and therefore loss of the dimethylarginine modification, is associated with a reduced specificity of CSR-1A for small RNAs targeting spermatogenesis genes and an increased binding of oogenic small RNAs.

## Discussion

The Argonaute protein CSR-1 is critical to promote germline gene expression and fertility. Here we demonstrate that, in addition to the short isoform CSR-1B which is expressed throughout germline development and during gametogenesis of both sexes, a longer isoform, CSR-1A, is selectively expressed during spermatogenesis. During L4, where both isoforms are expressed in the spermatogenesis region, CSR-1A and CSR-1B co-localize at P granules. Yet despite localizing to the same subcellular structure and sharing nearly complete sequence and presumably structural identity, CSR-1A and CSR-1B associate with distinct subsets of small RNAs targeting germline expressed genes – CSR-1A preferentially binds to small RNAs targeting spermatogenic genes, whereas CSR-1B binds to small RNAs targeting oogenic genes. The single exon unique to CSR-1A contains RG motifs that are modified with dimethylarginine and these methylated RG motifs are critical to CSR-1A specificity for spermatogenic small RNA substrates.

### CSR-1A promotes spermatogenic gene expression

Previous work has shown that CSR-1 can license targeted transcripts for expression in the adult germline, while its catalytic domain is necessary to tune embryonic gene expression and clear maternal transcripts

(Gerson-Gurwitz et al., 2016; Quarato et al., 2020; Seth et al., 2013; Wedeles et al., 2013). Though these studies did not make a distinction between CSR-1A and CSR-1B, based on the expression of only CSR-1B in the adult germline and embryos, we can attribute these CSR-1 activities to the CSR-1B isoform. Here we show that CSR-1A similarly does not down-regulate its target genes during spermatogenesis and in fact, seems to modestly promote spermatogenic gene expression.

The Argonaute proteins ALG-3 and ALG-4 similarly promote the expression of many spermatogenic target genes, and trigger the biogenesis of ALG-3/4-dependent 22G-RNAs as part of a feed-forward loop to promote paternal inheritance (Conine et al., 2013; 2010). With ALG-3 and CSR-1A sharing nearly identical expression patterns and CSR-1A being a spermatogenic Argonaute with a preference for spermatogenic 22G-RNAs, an obvious hypothesis would be that CSR-1A acts downstream of ALG-3 and ALG-4. However, the CSR-1A spermatogenic 22G-RNAs are produced independently of ALG-3/4 indicating that CSR-1A is likely part of a distinct pathway that targets the same spermatogenic genes as ALG-3/4 and another still-unidentified Argonaute binds the ALG-3/4-dependent 22G-RNAs. Importantly, these findings do not exclude the possibility of CSR-1B being an essential player in paternal epigenetic inheritance and acting downstream of ALG-3/4, as CSR-1B is still present in secondary spermatocytes with ALG-3, when CSR-1A has already disappeared (Conine et al., 2013).

If CSR-1A does not act downstream of ALG-3/4 for paternal inheritance, then how might it function to promote spermatogenic gene expression? If we look to CSR-1B as a guide, then perhaps CSR-1A acts to license spermatogenic transcripts for expression only in the L4 and male germlines by protecting them from targeting by piRNAs and the WAGO clade of Argonaute proteins. CSR-1A could further tune spermatogenic gene expression or clear spermatogenic transcripts at the spermatogenesis to oogenesis transition. Alternatively, the role for CSR-1A may be to sequester the abundant sperm transcripts and spermatogenic small RNAs away from CSR-1B, so that the dominant isoform can appropriately bind to its oogenic small RNA targets. Through this lens, CSR-1A serves almost exclusively to titrate CSR-1B siRNA levels targeting oogenic genes, and might explain the modest effect on the sperm transcripts upon removing CSR-1A. Further experiments will be needed to sort out these possibilities.

### Dimethylarginine promotes isoform-specific small RNA loading

Post-translational modification has been shown to play a key role for Argonaute function in *C. elegans* and in other systems. The *C. elegans* miRNA Argonaute protein, ALG-1, contains a cluster of phosphorylation sites that is conserved in human Argonaute Ago2, demonstrating that these modification sites are highly conserved between species. The function of this phosphorylation also appears to be conserved – the phosphorylated Argonaute proteins cannot associate with mRNAs, suggesting the role of the modification may be in mediating release of mRNA-Argonaute complexes for recycling (Quévillon Huberdeau et al., 2017). PIWI proteins from mouse, *Xenopus laevis*, and *Drosophila melanogaster* contain symmetrical dimethylarginines (sDMA), and in *Drosophila* this modification is mediated by protein arginine methyltransferase PRMT5 (Kirino et al., 2009; Reuter et al., 2009; Vagin et al., 2009). The dimethylarginine modification allows the Argonaute protein to interact with members of the Tudor domain protein family (Liu et al., 2010; Nishida et al., 2009; Reuter et al., 2009; Vagin et al., 2009; Webster et al., 2015). Tudor domains, which are protein-protein interaction modules, recognize methylated arginines and thus can mediate protein-protein interactions in a methylation-specific manner. Furthermore, for some PIWI proteins, subcellular localization to nuage is dependent on its interaction with Tudor proteins (Reuter et al., 2009; Vagin et al., 2009; Webster et al., 2015).

Here, we have found the first exon of CSR-1A to be highly modified with dimethylarginine. Interestingly, dimethylarginine is not required for CSR-1A localization to the P granule, but instead for small RNA specificity of CSR-1A, and thus recognition of the correct target transcripts. We hypothesize that CSR-1A interacts with an unknown Tudor domain protein through its dimethylated RG motifs, to promote engagement with a distinct small RNA biogenesis complex. In this scenario, dimethylarginine provides a new binding platform for proteins that cannot associate with CSR-1B, allowing CSR-1A to make distinct protein-protein interactions and ultimately to target a unique subset of genes. It is interesting to note that the majority of the RG motifs, including all methylated sites captured in our mass spectrometry experiment, are found in the exon unique to CSR-1A and absent from CSR-1B, making this highly modified region isoform-specific. There are several other *C. elegans* Argonaute proteins with multiple splice variants, including the miRNA Argonaute proteins ALG-1 and ALG-2, the oogenesis-specific primary Argonaute protein ERGO-1, and the WAGO-clade Argonaute protein PPW-1. It is currently unknown whether the isoforms of any of these genes have distinct functions or unique protein modifications, or if CSR-1 is unique in this regard.

## Supporting information

Key Resources Table

## Acknowledgements

We thank the members of the Phillips lab for helpful discussions and feedback on the manuscript. This work was supported by the National Institute of Health grants K22 CA177897 (to CMP), R35 GM119656 (to CMP), and T32 GM118289 (to DHN). CMP is a Pew Scholar in the Biomedical Sciences supported by the Pew Charitable Trusts (www.pewtrusts.org). Some strains were provided by the CGC, which is funded by NIH Office of Research Infrastructure Programs (P40 OD010440). Next generation sequencing was performed by the USC Molecular Genomics Core, which is supported by award number P30 CA014089 from the National Cancer Institute.

## Author Contributions

Conceptualization, DHN and CMP; Investigation, DHN; Formal Analysis – DHN and CMP; Writing – Original Draft, DHN and CMP; Writing – Reviewing & Editing, DHN and CMP; Supervision – CMP; Funding Acquisition – DHN and CMP

## STAR Methods

### Key Resources Table

#### Lead Contact and Materials Availability

Further information and requests for resources and reagents should be directed to, and will be fulfilled by the Lead Contact, Carolyn M. Phillips.

#### Experimental Model and Subject Details

Strains were maintained at 20°C on NGM plates seeded with OP50 *E. coli* according to standard conditions unless otherwise stated (Brenner, 1974). All strains used in this project are listed in the Key Resources Table.

## Methods Details

### Plasmid and strain construction

#### Plasmid-based CRISPR

All fluorescent and epitope tags were integrated at the endogenous loci by CRISPR genome editing (Arribere et al., 2014; Dickinson et al., 2015; 2013; Friedland et al., 2013; J. D. Ward, 2015). For all CRISPR insertions of fluorescent tags, we generated homologous repair templates using the primers listed in Table S3. Design of the *2xHA::mCherry* plasmid was described previously (Uebel et al., 2018). The *2xHA::mCherry[w/internal Floxed Cbr-unc-119(+)]* was amplified by PCR and assembled by isothermal cloning with ~1.5kb of sequence from either side of the *csr-1a* start codon (Gibson et al., 2009). *gfp::3xFLAG::wago-1*, *gfp::3xFLAG::csr-1a+b* and *gfp::3xFLAG::alg-3* were assembled into pDD282 (Addgene #66823) by isothermal assembly according to published protocols (Dickinson et al., 2015; Gibson et al., 2009). To protect the repair template from cleavage, we introduced silent mutations at the site of guide RNA targeting by incorporating these mutations into one of the homology arm primers or, if necessary, by performing site-directed mutagenesis (Dickinson et al., 2013). All guide RNA plasmids were generated by ligating oligos containing the guide RNA sequence into BsaI-digested pRB1017 (Addgene #59936) (Arribere et al., 2014). Guide RNA sequences are provided in Table S3. GFP/mcherry CRISPR injection mixes included 25-50 ng/μl repair template, 50 ng/μl guide RNA plasmid, 50 ng/μl *eft-3p::cas9-SV40_NLS::tbb-2 3’UTR* (Addgene #46168), 2.5–10 ng/μl GFP or mCherry co-injection markers. The *2xHA::mCherry::csr-1a* construct was injected into USC868 (*mut-16(cmp3[mut-16::gfp + loxP]) I; unc-119(ed3) III*), the *gfp::3xFLAG::wago-1* construct was injected into USC896 (*mut-16(cmp41[mut-16::mCherry::2xHA + loxP]) I*), the *gfp::3xFLAG::csr-1a+b* construct was injected into the wild-type strain, and the *gfp::3xFLAG::alg-3* construct was injected into both the wild-type strain and USC1066 (*csr-1a(cmp90[(2xHA + mCherry + loxP Cbr-unc-119(+) loxP)::csr-1a]) IV*) (Dickinson et al., 2015; 2013). For *csr-1a* deletions (*cmp135* and *cmp143*), *2xHA::csr-1a*, and *2xFLAG::csr-1a+b*, the injection mixes included 50 ng/μl repair oligo, 25 ng/μl guide RNA plasmid, 50 ng/μl pha-1 repair template, and *eft-3p::Cas9 + pha-1* guide (pJW1285, Addgene #61252). GE24 (*pha-1(e2123) III*) mutant animals were injected and subsequently shifted to restrictive temperature (25°C). Surviving F1 progeny were genotyped by PCR to identify the deletions and insertions of interest (J. D. Ward, 2015). For *2xFLAG::csr-1b*, the injection mix included 50 ng/μl *csr-1b* repair oligo, 25 ng/μl *csr-1b* guide RNA plasmid, 20 ng/μl *dpy-10* repair template, 25 ng/μl *dpy-10* guide RNA (pJA58, Addgene #59933), and 50 ng/μl *eft-3p::Cas9* (pJW1259, Addgene #61251). Mixture was injected into USC1065 (*csr-1a(cmp135) IV*) mutant animals. F1 animals with the Rol phenotype were isolated and genotyped by PCR to identify animals with the 2xFLAG insertion (Arribere et al., 2014). The *csr-1a* RG-to-AG mutants were created sequentially, starting with *csr-1a[7xAG]*. *csr-1a[4xAG]* was an incomplete repair event identified from the *csr-1a[7xAG]* injections. *csr-1a[11xAG]* was injected into the *csr-1a[7xAG]* mutant and *csr-1a[15xAG]* was subsequently injected into the *csr-1a[11xAG]* mutant. The *csr-1a RGG* injection mixes include 50 ng/μl repair template, 25 ng/μl each of two guide RNA plasmids, 25 ng/μl rol-6 guide RNA (pJA42, Addgene #59930), 20 ng/μl rol-6 repair template, and 50 ng/μl *eft-3p::Cas9* (pJW1259, Addgene #61251). F1 animals with the Rol phenotype were isolated and genotyped by PCR to identify animals with the RG-to-AG mutations (Arribere et al., 2014). All repair template sequences are provided in Table S3.

#### Protein-based CRISPR

For For *csr-1a* deletions (*cmp253* and *cmp254*), *csr-1b(cmp258)*, and *2xHA::csr-1a* in the *alg-3; alg-4* mutant, we used an oligo repair template and RNA guide (Table S3). The injection mixes for *csr-1b(cmp258)* and *2xHA::csr-1a* included 0.25 μg/μl Cas9 protein (IDT), 100 ng/μl tracrRNA (IDT), 14 ng/μl *dpy-10* crRNA, 42 ng/μl gene-specific crRNA, and 110 ng/μl of each repair template, and were injected into USC1066 (*csr-1a(cmp90[(2xHA + mCherry + loxP Cbr-unc-119(+) loxP)::csr-1a]) IV*) and WM200 (*alg-4(ok1041) III; alg-3(tm1155) IV*), respectively. The injection mix for *csr-1a(cmp253)* and *csr-1a(cmp254)* included 0.25 μg/μl Cas9 protein (IDT), 100 ng/μl tracrRNA (IDT), 14 ng/μl *dpy-10* crRNA, 21 ng/μl each gene-specific crRNA, and 110 ng/μl of each repair template, and was injected into the wild-type strain. The repair template was designed to create the *cmp254* mutation; the larger *cmp253* deletion was an incorrect repair event. Following injection, F1 animals with the Rol phenotype were isolated and genotyped by PCR to identify animals with the insertions and deletions of interest (Dokshin et al., 2018; Paix et al., 2015).

#### MosSCI

*csr-1* mCherry and GFP promoter fusions were integrated by Mos-mediated single-copy transgene insertion (MosSCI) (Frøkjaer-Jensen et al., 2008). For the MosSCI insertions of promoter-fused GFP and mCherry, we amplified *csr-1a* and *csr-1b* endogenous promoters, the mCherry and GFP genes, and the *csr-1* 3’UTR using the primers listed in Table S3. Plasmids were assembled by isothermal cloning (Gibson et al., 2009). For MosSCI injections, we integrated transgenes into the *ttTi5605 MosI* site in strain EG4322 (Ch. II) following a published MosSCI protocol (Frøkjaer-Jensen et al., 2008). Injection mixes contained 50 ng/μl MosSCI-targeting vector, 50 ng/μl *eft-3p*::*Mos1 transposase* (pCFJ601, Addgene #34874), 10 ng/μl *rab-3p*::*mCherry* (pGH8, Addgene #19359), 2.5 ng/μl *myo-2p*::*mCherry* (pCFJ90, Addgene #19327), 5 ng/μl *myo-3p*::*mCherry* (pCFJ104, Addgene #19328), and 10 ng/μl *hsp-16*.*1*::*peel-1*negative selection (pMA122, Addgene #34873) (Frøkjaer-Jensen et al., 2008).

### Antibody staining and imaging

Live imaging of *C*. *elegans* was performed in M9 buffer containing sodium azide to prevent movement. For immunofluorescence, *C. elegans* were dissected in egg buffer containing 0.1% Tween-20 and fixed in 1% formaldehyde in egg buffer as described (Phillips et al., 2009). Samples were immunostained with mouse anti-FLAG 1:500 (Sigma Aldrich, F1804), mouse anti-PGL-1 1:100 (DSHB K76) (Strome and Wood, 1983), and rat anti-HA 1:500 (Roche, 11867423001). Alexa-Fluor secondary antibodies were purchased from Thermo Fisher. Animals were dissected at the L4 (48 hours post hatching) or young adult stage (52 hours post hatching). Imaging was performed on a DeltaVision Elite microscope (GE Healthcare) using a 60x N.A. 1.42 oil-immersion objective. When data stacks were collected, three-dimensional images are presented as maximum intensity projections. Images were pseudocolored using the SoftWoRx package or Adobe Photoshop.

### Western blots

*C. elegans* were synchronized at 20°C by bleaching gravid adult animals and maintaining starved L1 larvae for at least 24 hours before plating on OP50. For sample collection, animals were harvested after 2 hours (L1), 12 hours (L2), 32 hours (L3), 50 hours (L4), and 72 hours (gravid adult) on OP50. Number of animals loaded per lane was normalized for actin – approximately 1000 L1s, 800 L2s, 600 L3s, 400 L4s, and 200 gravid adults. Proteins were resolved on 4-12% Bis-Tris polyacrylamide gels (Thermo Fisher), transferred to nitrocellulose membranes (Thermo Fisher), and probed with rat anti-HA-peroxidase 1:1,000 (Roche 12013819001), mouse anti-FLAG 1:1,000 (Sigma, F1804), mouse anti-actin 1:10,000 (Abcam ab3280), or rabbit anti-CSR-1 1:2,000 antibodies (Claycomb et al., 2009). Secondary HRP antibodies were purchased from Thermo Fisher.

### RNA Isolation and qRT-PCR

Synchronized animals were collected at L1, L2, L3, L4, and gravid adult stages, after two, 12, 32, 48, and 68 hours of feeding following L1 arrest, respectively. For mRNA libraries, synchronized L4 stage animals (~48 hours at 20°C and ~34 hours at 25°C after L1 arrest) were collected. RNA was isolated using Trizol reagent (Thermo Fisher), followed by chloroform extraction and isopropanol precipitation. RNA samples were normalized to 10μg/μL prior to DNase treatment (TURBO DNA-free kit, Thermo Fisher AM1907) and reverse transcribed with SuperScript III Reverse Transcriptase (Thermo Fisher 18080-051). All Real time PCR reactions were performed using the 2x iTaq Universal SYBR Green Supermix (Bio-Rad 1725121), following manufacturer’s protocols, and run in the CFX96 Touch Real-Time PCR System (Bio-Rad 1855195). Samples were normalized to *rpl-32* and Y45F10D.4 Primer sequences are available in Table S3.

### Brood size analysis

Wild-type and mutant *C. elegans* strains were maintained at 20**°**C prior to temperature-shift experiments. Animals were shifted to 25**°**C as L4 larvae and ten of their progeny were picked to individual plates for 25°C brood size analysis. To score the complete brood, each animal was moved to a fresh plate every day until egg-laying was complete. After allowing the progeny 2-3 days to develop, the total number of animals on each plate was counted.

### *In vitro* sperm activation assay

Virgin L4 males were isolated 24 hours before assay. 10-15 males were dissected in 30uL of 500μg/mL Pronase E (Millipore Sigma), which was dissolved in sperm medium (50 mM HEPES, 50 mM NaCl, 25 mM KCl, 5 mM CaCl2, 1 mM MgSO4, 1 mg/ml BSA), as described previously (S. Ward et al., 1983; Yen et al., 2020). Spermatids were incubated for 15 minutes at room temperature in a humid chamber before mounting and imaging on a DeltaVision Elite microscope (GE Healthcare) using a 60x N.A. 1.42 oil-immersion objective.

### Male fertility assay

Virgin L4 males were isolated 24 hours before assay and incubated at 20°C. L4 *unc-3* hermaphrodites were also selected 24 hours before assay and incubated at 20°C to allow for the first pass of egg-laying. On the day of the assay, four males were crossed to one hermaphrodite, 10 crosses per genotype. Crosses were incubated at 25°C and transferred to fresh plates every day for three days. Progeny were scored when they reached L3-L4 stage and the Unc phenotype was observable. Unc progeny indicated hermaphrodite sperm and wild-type progeny indicated male sperm (S. Ward and Carrel, 1979).

### Immunoprecipitations and mass spectrometry

For immunoprecipitation experiments followed by small RNA library preparation, ~100,000 synchronized L4 animals (~49 hours at 20°C after L1 arrest) or adult animals ~68 hours at 20°C after L1 arrest) were collected in IP Buffer (50 mM Tris-Cl pH 7.4, 100 mM KCl, 2.5 mM MgCl2, 0.1% Igapal CA-630, 0.5 mM PMSF, cOmplete Protease Inhibitor Cocktail (Roche 04693159001), and RNaseOUT Ribonuclease Inhibitor (Thermo Fisher 10777019)), frozen in liquid nitrogen, and homogenized using a mortar and pestle. After further dilution into IP buffer (1:10 packed worms:buffer), insoluble particulate was removed by centrifugation and 10% of sample was taken as “input.” The remaining lysate was used for the immunoprecipitation. Immunoprecipitation was performed at 4°C for 1 hour with pre-conjugated anti-HA affinity matrix (Roche 11815016001) or anti-FLAG affinity matrix (Sigma Aldrich A22220), then washed at least 3 times in immunoprecipitation buffer. A fraction of each sample was analyzed by western blot to confirm efficacy of immunoprecipitation. Trizol reagent (Thermo Fisher) was added to the remainder of each sample, followed by chloroform extraction, isopropanol precipitation, and small RNA library preparation.

For mass spectrometry experiments to identify post-translational modifications, immunoprecipitation was performed as described above, starting with ~1.25 million synchronized USC1110 (*csr-1a(cmp165[2xHA::csr-1a])*) L4 stage animals (~48 hours at 20°C after L1 arrest). Wild-type animals were prepped alongside as a negative control. Immunoprecipitation was performed using anti-HA affinity matrix (Roche 11815016001). After immunoprecipitation, a fraction of each sample was analyzed by western blot to confirm efficacy of immunoprecipitation. 2x sample buffer was added to the remainder of each sample, followed by gel electrophoresis (4-12% Bis-Tris polyacrylamide gels, Thermo Fisher) and overnight colloidal Coomassie staining. Bands containing immunoprecipitated protein were excised from gel and submitted to the Taplin Mass Spectrometry facility at Harvard Medical school for identification of post-translational modifications.

### Small and mRNA library preparation

Small RNAs (18 to 30-nt) were size selected on denaturing 15% polyacrylamide gels (Bio-Rad 3450091) from total RNA samples. Small RNAs were treated with 5’ RNA polyphosphatase (Epicentre RP8092H) and ligated to 3’ pre-adenylated adapter with Truncated T4 RNA ligase (NEB M0373L). Small RNAs were then hybridized to the reverse transcription primer, ligated to the 5’ adapter with T4 RNA ligase (NEB M0204L), and reverse transcribed with Superscript III (Thermo Fisher 18080-051). Small RNA libraries were amplified using Q5 High-Fidelity DNA polymerase (NEB M0491L) and size selected on a 10% polyacrylamide gel (Bio-Rad 3450051).

Library concentration was determined using the Qubit 1X dsDNA HS Assay kit (Thermo Fisher Q33231) and quality was assessed using the Agilent BioAnalyzer. Libraries were sequenced on the Illumina NextSeq500 (SE 75-bp reads) platform. For mRNA sequencing, total RNA samples were submitted in triplicate to Novogene Genome Sequencing Company for library preparation. Libraries were sequenced on the Illumina (PE 150-bp reads) platform.

### Clustal Omega alignments and IUPred Disorder prediction

Clustal Omega alignment was performed using protein sequences for CSR-1 orthologs available on Wormbase (www.wormbase.org) (Sievers et al., 2011). When more than one CSR-1 ortholog was present in a given species, a single protein sequence was selected for analysis. Proteins are *C. brenneri* CBN29996 (WBGene00191961), *C. briggsae* CSR-1 (WBGene00037276), *C. elegans* CSR-1, isoform a (WBGene00017641), *C. japonica* CSR-1 (WBGene00126657), *C. latens* PRJNA248912_FL83_15994, *C. nigoni* CSR-1 (PRJNA384657_Cni-csr-1), *C. sinica* PRJNA194557_Csp5_scaffold_00781.g15416.t2, *C. remanei* PRJNA248911_FL82_23103, *C. tropicalis* PRJNA53597_Csp11.Scaffold629.g12789.t1. For *C. japonica*, exon 1 of CSR-1A was manually annotated from the CJA07453.1 transcript. For disorder prediction, we used IUPred2A (https://iupred2a.elte.hu/) with long disorder parameters and the same protein sequences as were used for Clustal Omega alignment (Dosztányi et al., 2005; Mészáros et al., 2018).

### Quantification and Statistical Analysis

#### Bioinformatic Analysis

For small RNA libraries, sequences were parsed from adapters using FASTQ/A Clipper (options: -Q33 -l 17 -c -n -a TGGAATTCTCGGGTGCCAAGG) and quality filtered using the FASTQ Quality Filter (options: -Q33 -q 27 -p 65) from the FASTX-Toolkit (http://hannonlab.cshl.edu/fastx_toolkit/), mapped to the *C. elegans* genome WS258 using Bowtie2 v. 2.2.2 (default parameters) (Langmead and Salzberg, 2012), and reads were assigned to genomic features using FeatureCounts (options: -t exon -g gene_id -O --fraction –largestOverlap) which is part of the Subread v. 1.5.1 package (Liao et al., 2014; 2013). Differential expression analysis was done using DESeq2 v. 1.22.2 (Love et al., 2014). To define gene lists from IP experiments, a 2-fold-change cutoff, a DESeq2 adjusted p-value of ≤0.05, and at least 10 RPM in the IP libraries were required to identify genes with significant changes in small RNA levels. Additionally, any genes identified as having differentially enriched small RNAs from control samples (HA or FLAG immunoprecipitations from wild-type animals), were removed from further analysis.

For mRNA libraries, sequences were parsed from adapters using Trimmomatic v. 0.36 (options: PE -phred33 ILLUMINACLIP:<fasta with adaptor sequences>:2:30:10 LEADING:3 TRAILING:3 SLIDINGWINDOW:4:30 MINLEN:30) (Bolger et al., 2014) and mapped to the *C. elegans* genome WS258 using HISAT2 v. 2.1.0 (options: --dta-cufflinks --known-splicesite-infile <path to file of known splice sites>) (Kim et al., 2015). Differential expression analysis was done using Cuffdiff v. 2.1.1 (default parameters) (Trapnell et al., 2012; 2010).

CSR-1 target genes, male CSR-1 target genes, ALG-3/4 target genes, *mutator* target genes, germline-enriched genes, spermatogenesis-enriched genes, and oogenesis-enriched genes were previously described (Conine et al., 2013; Gu et al., 2009; Lee et al., 2012; Phillips et al., 2014; Reinke et al., 2004; Rogers and Phillips, 2020; Tsai et al., 2015; Zhang et al., 2011). Additional data analysis was done using R, Excel, and Python. Venn diagrams were generated using BioVenn (Hulsen et al., 2008) and modified in Adobe Illustrator. Sequencing data is summarized in Table S4.

## Data and Code Availability

High-throughput sequencing data for RNA-sequencing libraries generated during this study are available through Gene Expression Omnibus (GSE151828).

## Supplemental Figures

**Figure S1.**
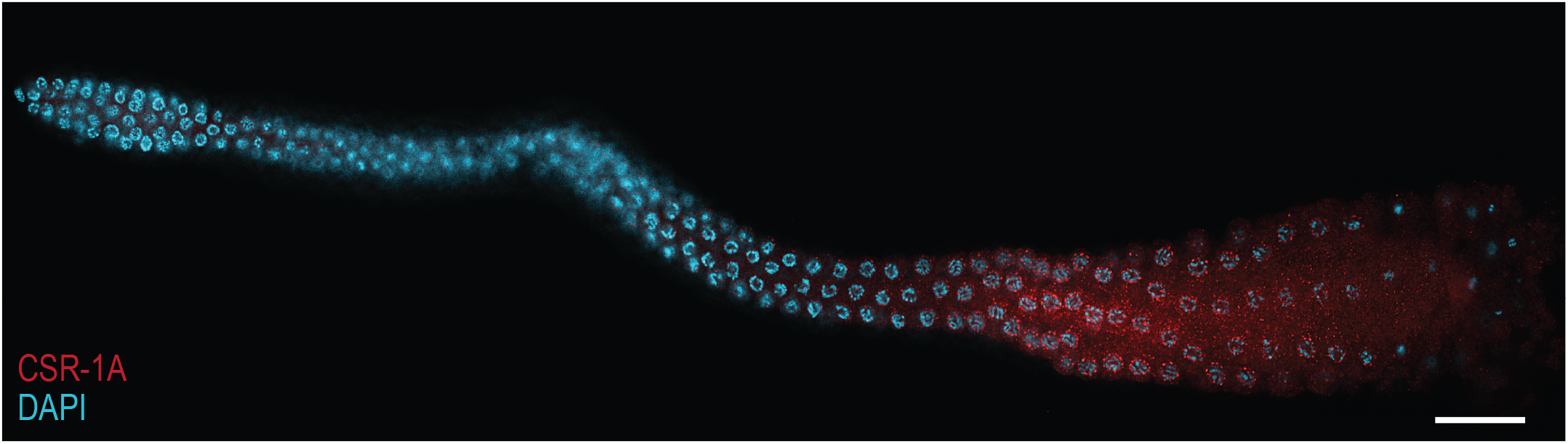
CSR-1A is expressed in the spermatogenesis region of the male germline. Immunofluorescence staining of CSR-1A in a dissected L4 male germline carrying the 2xHA::mCherry::CSR-1A transgene. HA antibodies are used to recognize CSR-1A and DNA is stained with DAPI. Scale bar, 20μM.

**Figure S2.**
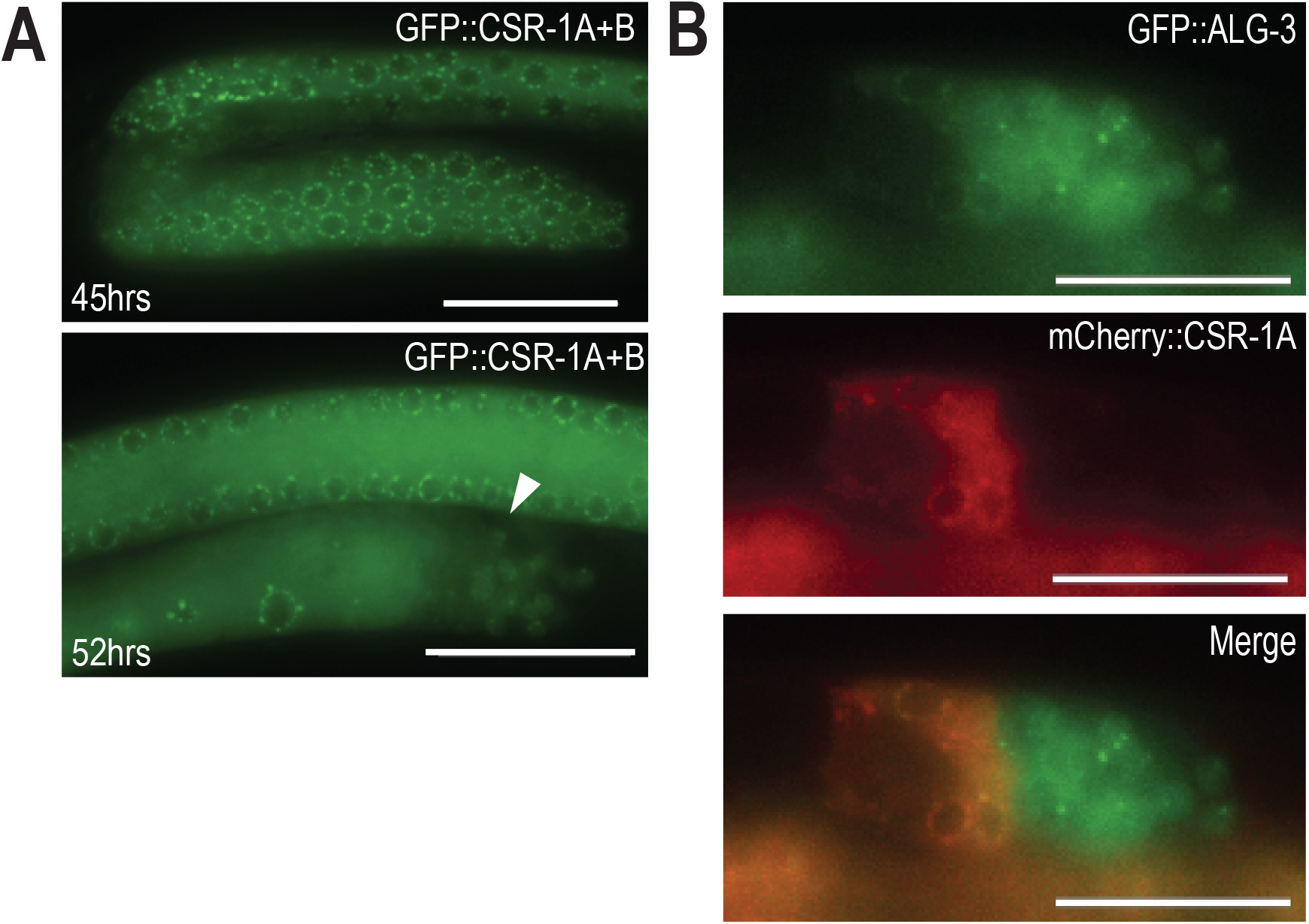
CSR-1A is excluded from secondary spermatocytes. (A) 3xFLAG/GFP::CSR-1A+B at 45 hrs (early L4 stage) or 52 hrs (young adult stage) post-L1 arrest. White arrows indicate where region of secondary spermatocytes. (B) Live imaging of double-transgenic animal labelled for both GFP::ALG-3 and mCherry::CSR-1A at the young adult stage. Scale bars, 25μM.

**Figure S3.**
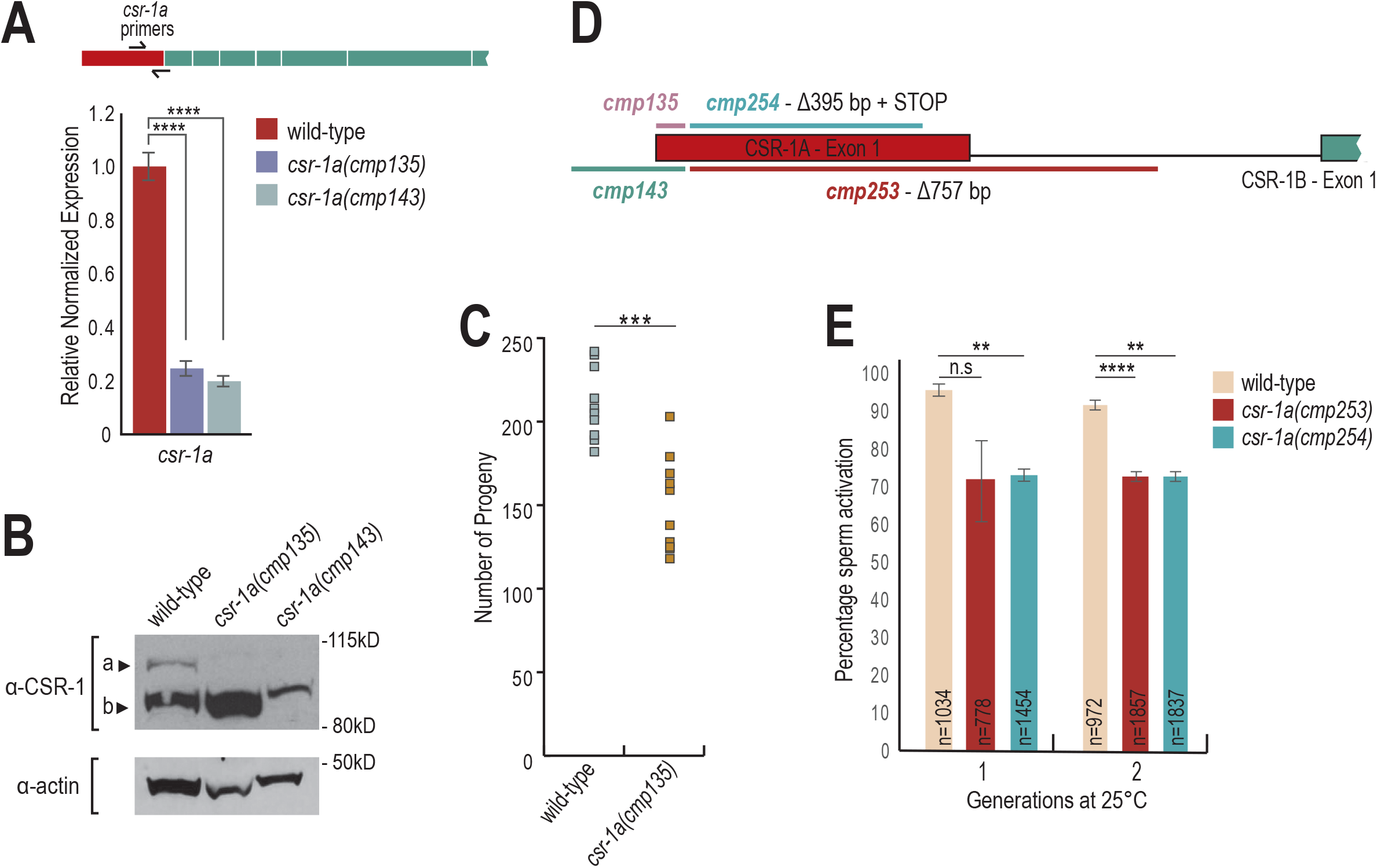
CSR-1A is required for optimal male fertility. (A) RT-qPCR on *csr-1a* mutants (*cmp135* and *cmp143*) and wild-type animals at L4. Relative expression was normalized over *rpl-32* and calculated relative to wild-type. **** indicates a p-value≤0.0001. (B) Western blot detecting for both CSR-1 isoforms expression in wild-type and *csr-1a* mutant L4 hermaphrodites, using CSR-1 antibody. Actin is shown as a loading control. (C) Brood size assay on *csr-1a* (*cmp135*) raised at 25°C (n=10). Two-tail t-tests were performed to determined statistical significance. *** indicates a p-value≤0.001 (D) Schematic representation of two new *csr-1a* mutants (*cmp253* and *cmp254*) (E) *in vitro* sperm activation assay on additional *csr-1a* alleles and wild-type. Animals were raised at 25°C for either one or two generations. Each experiment was performed in triplicate. At least 100 spermatids were counted for each replicate for each condition, for a total of at least 400 spermatids per condition. Two-tail T-tests were performed to determine statistical significance. ** indicates a p-value≤0.01 and **** indicates a p-value≤0.0001.

**Figure S4.**
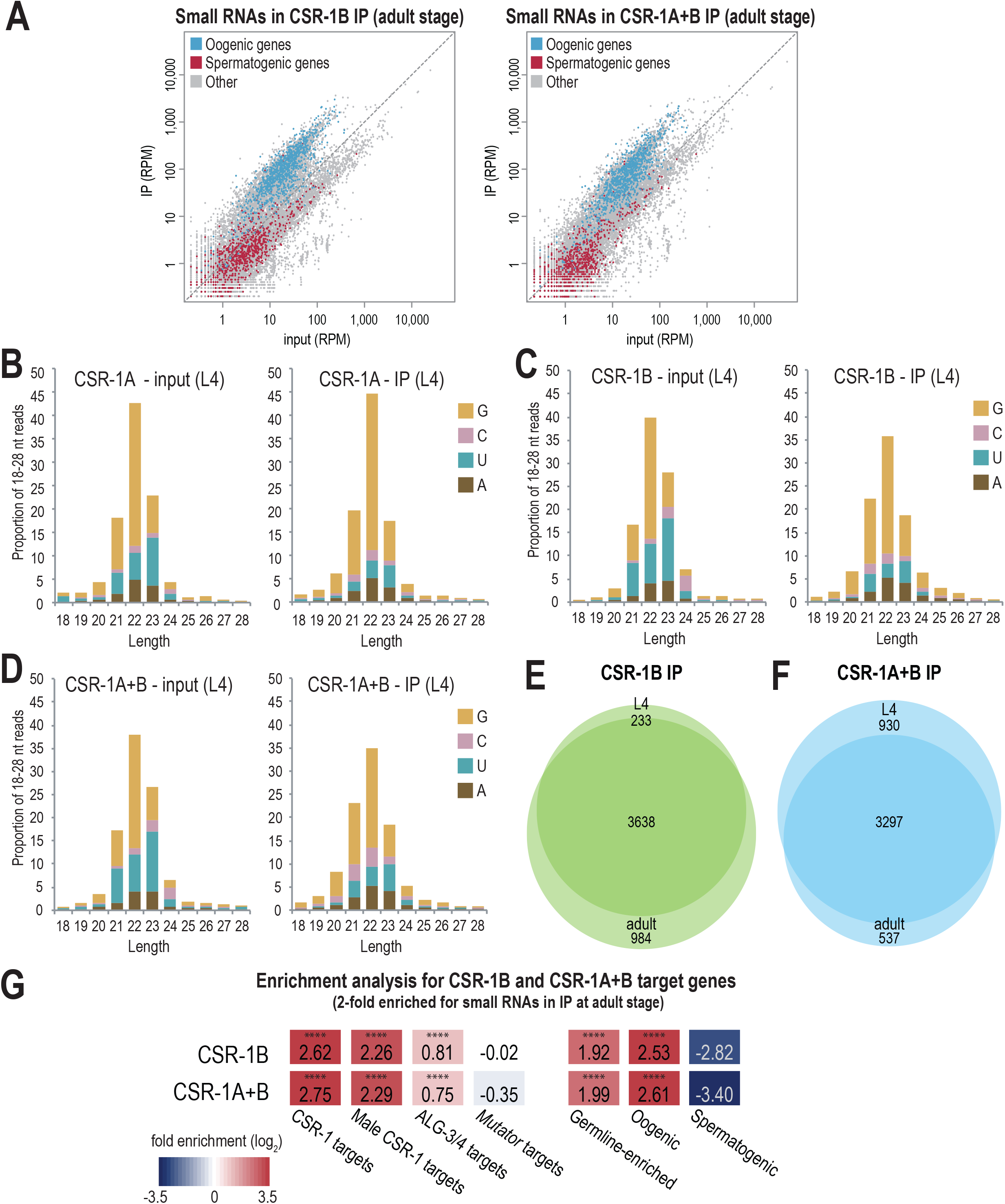
Both CSR-1 isoforms bind germline-enriched 22G-RNAs. (A) Normalized reads for oogenesis and spermatogenesis genes at adult stage from FLAG::CSR-1B IP and FLAG::CSR-1A+B IP compared to input. are Oogenesis and spermatogenesis genes are indicated in blue and red respectively. (B-D) 5’ length and nucleotide distribution in representative input and IP libraries from L4 stage HA::CSR-1A (B), FLAG::CSR-1B (C), and FLAG::CSR-1A+B (D). (E-F) Venn diagram indicates overlap of L4 and adult target genes from CSR-1B (E) and CSR-1A+B (F) immunoprecipitations. (G) Enrichment analysis (log2(fold enrichment)) examining the overlap of adult stage CSR-1B and CSR-1A+B target genes with known targets of the CSR-1, male CSR-1, ALG-3/4, and *mutator* small RNA pathways and oogenesis and spermatogenesis-enriched genes. See Materials and Methods for gene list information. Statistical significance for enrichment was calculated using the Fisher’s Exact Test function in R. **** indicates a p-value≤0.0001.

**Figure S5.**
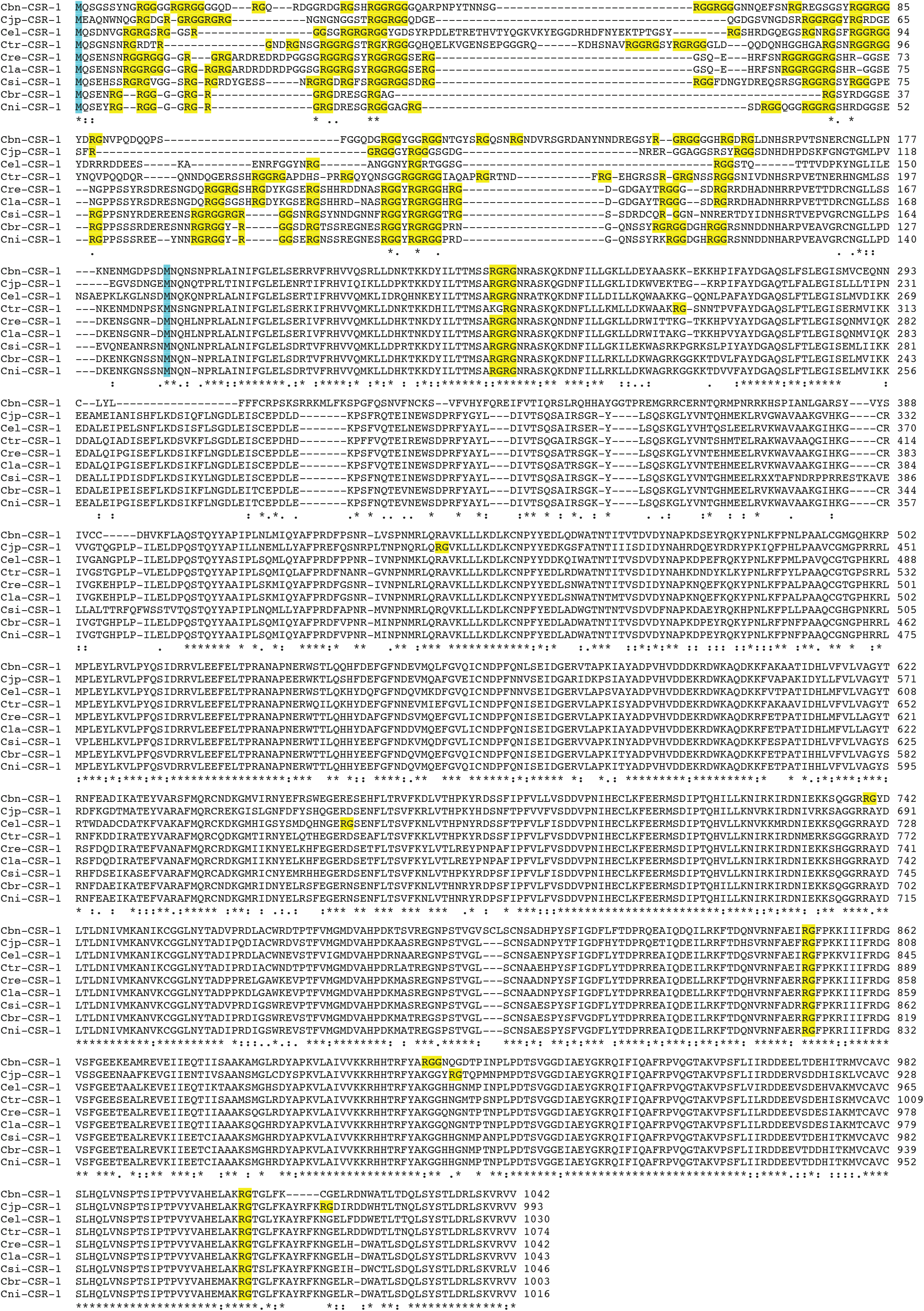
Clustal Omega alignment of CSR-1 in Caenorhabditis species. Arginine/glycine motifs are highlighted in yellow and start codons for both CSR-1A and CSR-1B are highlighted in blue. See Materials and Methods for protein sequence information.

**Figure S6.**
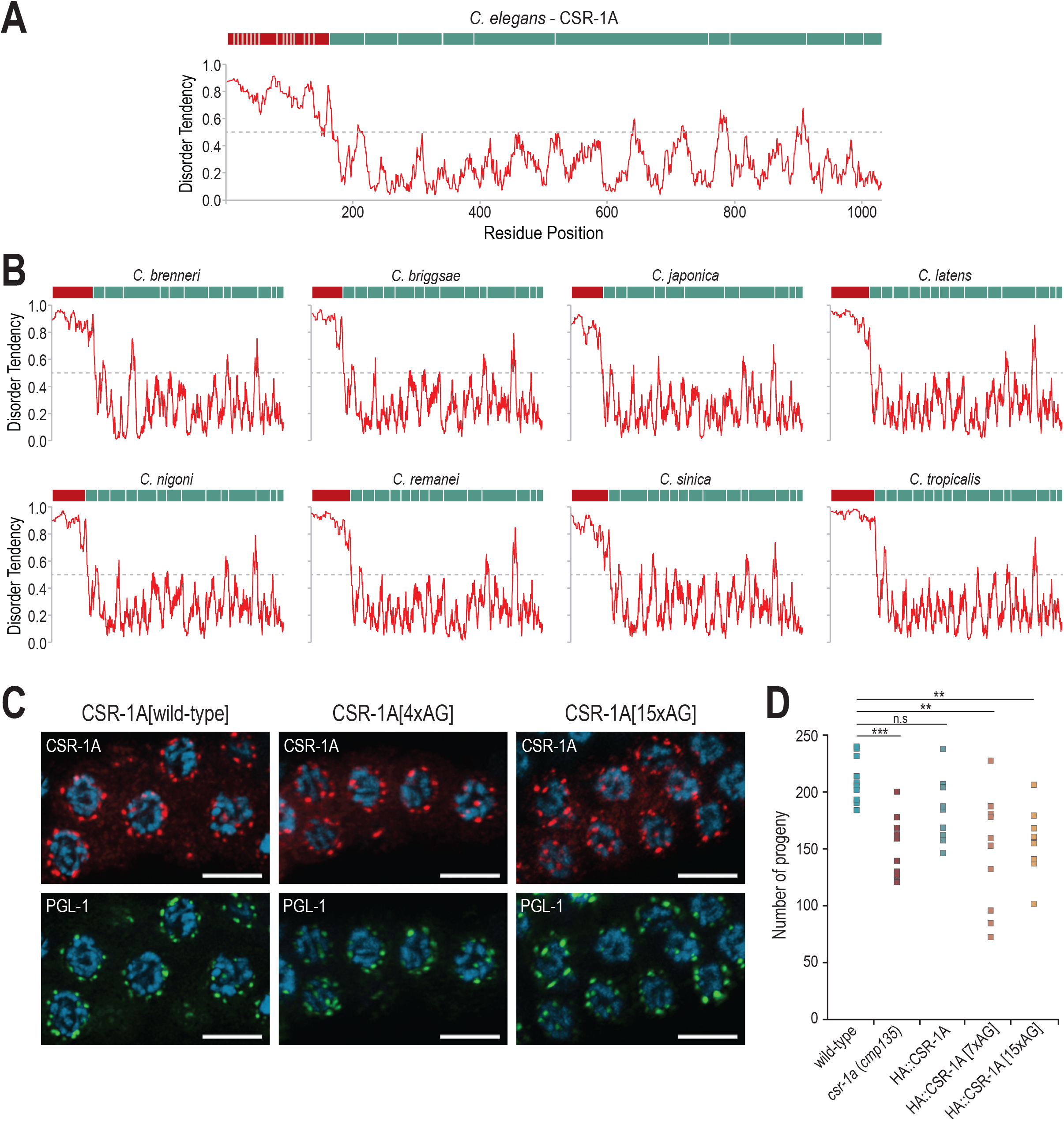
The N-terminal exon of CSR-1 is disordered across Caenorhabditis species. (A-B) Graphs examining the disorder tendencies of CSR-1A in *C. elegans* (A) and CSR-1 in related nematode species (B). Residue position is plotted on the x-axes and disorder tendency scores (y-axes) above 0.5 indicate disorder. Red bar above each plot marks region of first exon and green bars mark all other exons. (C) Immunofluorescence staining of dissected L4 hermaphrodite germlines in HA::CSR-1A, HA::CSR-1A[4xAG], and HA::CSR-1A[15xAG], using antibodies against HA and PGL-1. Scale bars, 5μM. (D) Brood size assay on wild-type and mutant animals raised at 25°C (n=10). Two-tail T-tests were performed to determined statistical significance. ** indicates p-value≤0.01, *** indicates a p-value≤0.001.

**Table S1.** Small RNA enrichment in CSR-1 immunoprecipitations.

**Table S2.** Peptides captured from 2xHA::CSR-1A IP-mass spectrometry experiment.

**Table S3.** Oligonucleotides sequences used in this study.

**Table S4.** Sequencing library statistics.

